# The gene regulatory landscape driving mouse gonadal supporting cell differentiation

**DOI:** 10.1101/2024.12.09.627451

**Authors:** Isabelle Stévant, Elisheva Abberbock, Meshi Ridnik, Roni Weiss, Linoy Swisa, Christopher R Futtner, Danielle M Maatouk, Robin Lovell-Badge, Valeriya Malysheva, Nitzan Gonen

## Abstract

Gonadal sex determination relies on tipping a delicate balance involving the activation and repression of several transcription factors and signalling pathways. This is likely mediated by numerous non-coding regulatory elements that shape sex-specific transcriptomic programs. To explore the dynamics of these in detail, we performed paired time-series of transcriptomic and chromatin accessibility assays on pre-granulosa and Sertoli cells throughout their development in the embryo, making use of new and existing mouse reporter lines. Regulatory elements were associated with their putative target genes by linkage analysis, and this was complemented and verified experimentally using promoter capture Hi-C. We identified the transcription factor motifs enriched in these regulatory elements along with their occupancy, pinpointing LHX9/EMX2 as potentially critical regulators of ovarian development. Variations in the DNA sequence of these regulatory elements are likely to be responsible for many of the unexplained cases of individuals with Differences of Sex Development.

**Teaser:** Multiomics analysis revealed the regulatory elements and transcription factors responsible for gonadal sex determination.

## Introduction

In eutherian mammals, sex determination occurs during embryo development with the bipotential gonad committing to either testicular or ovarian cell fates (*1*). This choice relies on an initially extremely delicate balance between the expression and repression of several key transcription factors (TFs) and signalling pathways. There is then a period of reinforcement involving redundant mechanisms, perhaps to ensure the whole gonad follows one fate, before specific factors can become predominant postnatally (*2*, *3*). The gonadal differentiation process is, therefore, an ideal one in which to study cell fate decisions and the roles played by gene regulatory networks over time (*4*).

In mice, the bipotential gonad begins to develop at embryonic day 10 (E10.0) as a thickening layer positioned on the ventromedial surface of the mesonephros (*1*, *5–7*). The early bipotential gonad in the mouse comprises two distinct populations of progenitor cells: those derived from the coelomic epithelium, which can be considered multipotent because it is the origin of most of the somatic cell types, and the primordial germ cells, which differentiate according to the type of gonad that forms into spermatogonia in the testis, or oogonia in the ovary. The somatic cell types include the bipotential supporting cell precursors that differentiate into Sertoli cells (male) or pre-granulosa cells (female), and the steroidogenic cell precursors, which give rise to the majority of Leydig (male) or theca (female) cells (*1*, *6*– *10*). The underlying mesonephros also contributes to the gonadal via progenitor cells that later differentiate as either perivascular, Leydig and myoid cells (male) (*8*, *11*, *12*) and theca cells (female) (*13*). More recently, a population of supporting-like cells deriving from the gonadal coelomic epithelium close to the mesonephros have been identified at the origin of the *rete testis* and *ovarii* (*9*, *14*).

In XY gonads, the sex-determining gene *Sry*, located on the Y chromosome, together with its direct downstream gene *Sox9*, initiates a genetic cascade at E11.5 leading to the differentiation of the supporting cell precursors into Sertoli cells (*15–21*). Sertoli cell identity is further maintained by several other TFs and signalling pathways including NR5A1, GATA4, WT1, DMRT1, SOX8, and FGF9, which together reinforce Sertoli cell fate and repress the pre-granulosa cell pathway (*1*, *6*, *7*). Once established, Sertoli cells instruct the steroidogenic cell precursors to differentiate into Leydig cells (*22*), and also encapsulate the germ cells and promote their differentiation into prospermatogonia (*23*).

In XX gonads, without SRY to upregulate *Sox9* expression by E11.5, the activity of ovarian promoting factors leads to pre-granulosa cell differentiation and ovary development. These include the ovarian determining factor WT1^-KTS^ isoform, the WNT4/RSPO1/β-Catenin signalling pathway and several TFs, notably FOXL2 and RUNX1 (*24–31*). The expression of these factors within the supporting cell precursors also act to repress the Sertoli cell differentiation program, for example high levels of WT1^-KTS^ interfere with *Sry* expression (*26*), while WNT signalling interferes with *Sox9* upregulation (*27*, *32*).

Despite identifying many key pro-testicular and pro-ovarian factors, the gene regulatory networks they govern and the genomic elements they utilize to coordinate gene expression during sex determination remain poorly understood. *Cis*-regulatory elements are defined as regions of non-coding of DNA (usually 500–1,500 bp) that have the capacity to control gene expression. Such elements are highly enriched in transcription factor binding sites and can be located upstream or downstream and at varying distances of the transcriptional start site(s) of the genes they regulate (*33–35*). *Cis*-regulatory elements are usually classified as promoters, enhancers, silencers, and insulators. Within the field of sex determination, few *cis*-regulatory regions have been characterized in detail to date. We have previously identified Enh13, a distal enhancer of the *Sox9* gene, as a critical enhancer for male sex determination. We and others have shown that deletions, duplications and even micro-deletions in two transcription factor binding sites (TFBS) of this enhancer can significantly alter *Sox*9 expression levels, leading to complete sex reversal in both mice and humans (*36–39*). Thus, we believe Enh13 is one of many regulatory elements in which variant DNA sequences may underlie numerous unexplained cases of Differences of Sex Development (DSDs). This data will also offer insights into the complex gene regulatory networks driving sex determination and gonadal differentiation.

Pioneer attempts to characterize the gene regulatory landscape of purified Sertoli and pre-granulosa cells have used DNaseI-seq (*40*) and ATAC-seq (*41*) on TESCO-CFP (*21*) and TESMS-CFP (*37*) transgenic lines carrying fluorescent reporter constructs controlled by the native and a mutated version of the TESCO enhancer of *Sox9*, respectively. While these approaches identified key elements like Enh13 (*37*), they provided limited temporal resolution. Indeed, DNaseI-seq was restricted to Sertoli cells at E13.5 and E15.5, and ATAC-seq examined supporting cell precursors at E10.5 and Sertoli and pre-granulosa cells at E13.5. However, single-cell transcriptomics analyses indicate that critical gonadal gene expression changes occur between E11.5–E12.5 (*9*, *10*, *42*), likely accompanied by dynamic patterns of chromatin accessibility. The lack of such data between E10.5 and E13.5 may obscure transient regulatory elements active during this critical window. Thus, a detailed timeline of chromatin accessibility, supported by high-quality, cell type-specific, and time-resolved bulk RNA datasets, is essential to fully capture and begin to understand the complexity of *cis*-regulation during sex determination.

Here, we generated a novel reporter mouse strain, *Enh8-mCherry,* which allowed us to efficiently purify pre-granulosa cells. Using both the *Enh8*-mCherry and *Sox9^IRES-GFP^* mouse strains, we purified pre-granulosa and Sertoli cells at four developmental time points in which sex determination occurs and performed bulk RNA-seq and ATAC-seq. This data constitutes the first bulk RNA-seq data of these critical cell types and provides a wealth of information of their dynamic transcriptomic profiles. The chromatin accessibility data enables the identification of numerous putative regulatory elements that may function during sex determination, among which many exhibit sex- and stage-specific patterns. We employed computational approaches to provide linkage correlation between accessible regions and gene expression profiles allowing us to identify putative enhancers and silencers of key sex determination genes. This was validated *in vivo* by performing Promoter Capture Hi-C (PCHi-C) on purified Sertoli and pre-granulosa cells at E13.5. We find enrichment of our ATAC-seq candidate regulatory elements within the PCHi-C data, suggesting that many of the elements we identified are physically bound to the promoters of the neighbouring genes that they regulate within these cells. Furthermore, we performed transcription factor motif and footprinting analyses on the open chromatin regions in each sex, allowing us to identify the regulators that are bound to them and therefore likely to control the process of sex determination. In Sertoli cells, we see enriched binding of SOX/SRY/DMRT1/GATA/WT1 and NR5A1 TFs, all of which are known to be critical for testis development (*43*). Strikingly, in pre-granulosa cells we see all of the known pro-female factors as WT1/FOXL2/RUNX1/TCF and GATA, but the most enriched motif is one bound by EMX2 and LXH9, suggesting that these factors may play an additional, yet unidentified role during ovary differentiation and not only during early gonad development (*1*, *44*, *45*). Altogether, this study constitutes an extensive analysis of the gene regulatory networks that operate during mouse sex determination, and it is highly likely that variants within the human homologues of these regulatory elements are responsible for many of the undiagnosed cases of DSD.

## Results

### Generation and use of transgenic mouse lines that allow efficient purification of pre-granulosa and Sertoli cells

To explore the gene regulatory network and *cis*-regulatory elements controlling gonadal sex determination, we first needed to purify gonadal supporting cells from both sexes. Therefore, we needed transgenic mouse lines that allow for the efficient purification of pre-granulosa and Sertoli cells at key developmental stages. Existing pre-granulosa cell reporters, such as TESMS-CFP (*41*) or Sry-GFP (*46*, *47*) present limitations in term of percentage of positive cells and sorting efficiency due to low fluorescent signals. To overcome these limitations, we generated a novel transgenic mouse line in which the *Sox9* enhancer 8 (referred to as Enh8) was cloned upstream of the *hsp68* minimal promoter and the *mCherry* gene (**Fig. 1A**, termed *Tg(Enh8-mCherry),* or in short *Enh8-mCherry*). Enh8 is a 672 bp-long enhancer, located 838 kb upstream to the *Sox9* transcription start site (TSS) (*37*). This enhancer has been shown to be active and capable of driving LacZ reporter expression in the mouse embryonic ovary. Indeed, although *Sox9* is highly expressed in Sertoli cells and faintly in pre-granulosa cells, several studies performing ChIP-seq in both embryonic and adult ovaries have found binding of FOXL2 and RUNX1, two transcription factors involved in the maintenance of pre-granulosa cell identity, to *Sox9* Enh8 (**Fig. 1A**) (*28*, *48–51*). The binding of these factors explains why this enhancer is active in pre-granulosa cells. To characterize the expression profile of the *Enh8-mCherry* mouse line, we dissected XX and XY embryonic gonads at E11.5, 12.5, 13.5 and 15.5. As expected, mCherry expression was evident in ovaries from E11.5 onward (**Fig. 1B**). Surprisingly, and conversely to what we observed with the LacZ reporter (*37*), mCherry expression was also evident in E11.5-E15.5 testes (**Fig. S1A**).

**Figure 1.**
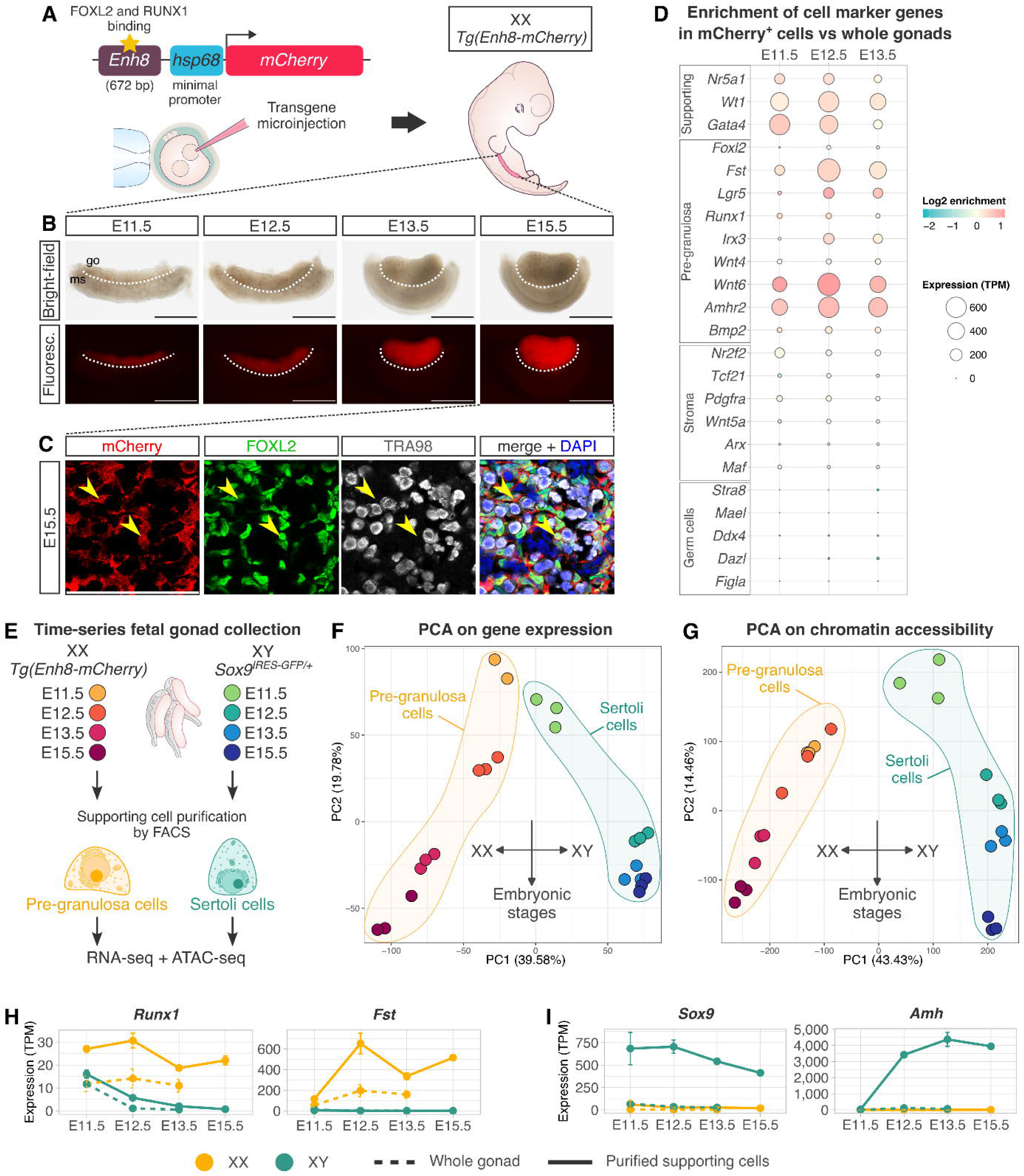
Generation of a new mouse line allowing pre-granulosa cell purification for multi-omics analysis. (**A**) Enh8 was cloned upstream of the mCherry reporter gene controlled by the *hsp68* minimal promoter. The obtained vector was injected into zygote mouse embryos by microinjection to obtain the *Enh8-mCherry (Tg(Enh8-mCherry))* mouse line. (**B**) Binocular pictures of dissected XX *Enh8-mCherry* gonads at E11.5, E12.5, E13.5 and E15.5 in bright-field and fluorescence. go=gonad, ms=mesonephros. Scale bar: 500 µm. (**C**) Immunofluorescent staining of E15.5 *Enh8-mCherry* ovary sections. mCherry (transgene) is labelled in red, FOXL2, a marker of pre-granulosa cells is labelled in green, and TRA98, a marker of germ cells is labelled in grey. Some mCherry expressing pre-granulosa cells are highlighted with yellow arrowheads. Scale bar: 100 µm. (**D**) Gene expression enrichment of ovarian marker genes in *Enh8-mCherry* sorted cells at E11.5, E12.5 and E13.5 RNA-seq compared to whole gonad RNA-seq (from Zhao et al. 2018 (*52*)). Enrichment (log2 of the ratio between *Enh8-mCherry* sorted cells and whole gonad gene expression) is represented with a blue-to-red gradient, and the expression levels of the *Enh8-mCherry* sorted cells are represented as TPM (transcript per million) with the size of the dots. (**E**) Schematic experimental design. XX gonads from *Enh8-mCherry* and XY gonads from *SOX*9^IRES-GFP/+^ mouse lines were collected at E11.5, E12.5, E13.5 and E15.5. Pre-granulosa and Sertoli cells were purified by FACS at each embryonic stage, and the sorted cells were subjected to RNA-seq and ATAC-seq to constitute a time-series paired gene expression and chromatin accessibility data collection. (**F**) and (**G**) PCA (principal component analysis) of the obtained RNA-seq and ATAC-seq data. Samples are coloured by sex and embryonic stage. Pre-granulosa samples were circled in yellow, and Sertoli cell samples in green. (**H**) and (**I**) Expression profiles of *Runx1*, *Fst* (pre-granulosa specific genes), and *Sox9* and *Amh* (Sertoli specific genes) in purified cells (continuous line) and in whole gonads (from Zhao et al. 2018 (*52*), dashed line) along embryonic stages in both sexes (XX in yellow, XY in green).

To explore whether the mCherry-positive cells are indeed pre-granulosa cells, we performed co-immunostaining with antibodies against mCherry, FOXL2 (pre-granulosa cells) and TRA98 (germ cells) in sections of ovaries. As demonstrated in **Fig. 1C** and **Fig**. **S1B**, the cytoplasmic mCherry staining overlaps well with nuclear FOXL2 staining suggesting that the mCherry labels pre-granulosa cells. No overlap is seen with the TRA98-positive germ cells or COUP-TFII (NR2F2)-positive stromal cells. To further confirm that the XX mCherry-positive cells are pre-granulosa cells, we performed bulk RNA-seq on E11.5, E12.5 and E13.5 mCherry-positive sorted cells. We compared the expression of different cell-type-specific genes (*9*, *10*) in our RNA-seq data to that of embryonic whole gonad RNA-seq data (*52*). As demonstrated in **Figure 1D**, expression profiles of the mCherry-positive cells resemble those of pre-granulosa cells with enriched expression of markers of the pre-supporting cells, but also many markers known to be expressed in pre-granulosa cells. No expression or negative enrichment was found with markers of both germ cells and stromal cells, indicating the absence of these cells in the mCherry-positive cell population (**Fig. 1D**). All of the above strongly suggest that this new *Enh8-mCherry* reporter mouse line is an efficient genetic tool to allow the sorting of pure populations of pre-granulosa cells from embryonic XX gonads.

To analyse which cells are mCherry-positive in embryonic testes, we performed immunostaining with antibodies against mCherry, SOX9 (Sertoli cells) and TRA98 (germ cells) (**Fig. S1C**). While mCherry overlapped with SOX9, suggesting it is labelling Sertoli cells, there was also expression outside the tubules, within the interstitium. Co-staining with mCherry and 3βHSD indicated that the reporter is also labelling fetal Leydig cells (**Fig. S1C**). Hence, this mouse line cannot be used to purify Sertoli cells from embryonic gonads. Instead, we used the well-established *Sox9^IRES-GFP^* reporter strain (*53*) where GFP labels *Sox9*-expressing Sertoli cells (**Fig. S1D**).

We and others have previously used the TESCO-CFP and TESMS-CFP reporter mouse lines to purify embryonic Sertoli and pre-granulosa cells, respectively (*37*, *40*, *41*). While these lines allow the purification of Sertoli and pre-granulosa cells, the percentage of CFP-positive cells out of the entire embryonic gonad was significantly lower than we find with the *Enh8-mCherry* or *Sox9^IRES-GFP^* lines (∼33-40% mCherry-positive cells in *Enh8-mCherry* ovaries, ∼14% of GFP-positive cells in *Sox9^IRES-GFP^* testes, ∼5-7% CFP positive cells with the TESCO-CFP/TESMS-CFP reporters) (**Fig. S2**). ScRNA-seq studies show that the actual proportion of supporting cells in the embryonic gonads is similar to those obtained after sorting with the *Enh8-mCherry* or the *Sox9^IRES-GFP^* lines (**Fig. S2D**), suggesting that these allow more representative capture of Sertoli and pre-granulosa cell populations at these stages.

### Exploring the transcriptomics and chromatin accessibility of purified pre-granulosa and Sertoli cells from embryonic gonads

To investigate the establishment of the *cis*-regulatory element landscape that drives pre-supporting cell differentiation into pre-granulosa and Sertoli cells, we performed paired time-series transcriptomic (RNA-seq) and chromatin accessibility assays (ATAC-seq) at four time points (E11.5, E12.5, E13.5 and E15.5) covering cell fate commitment and differentiation in both sexes using *Enh8-mCherry* for XX and *Sox9^IRES-GFP^* for XY gonads (**Fig. 1E**, **Fig. S3**, methods).

The bulk transcriptomes on sorted Sertoli and pre-granulosa cells provided the expression level of 12,058 protein-coding genes, and the ATAC-seq detected a total of 87,988 high confidence and non-overlapping open chromatin regions (stages and sexes combined), which represent 2.8% of the mouse genome. Principal component analysis (PCA) reveals that transcriptomic and chromatin accessibility changes along supporting cell differentiation are following fairly similar trajectories and are associated first with the sex (PC1), and second with the embryonic stages (PC2) (**Fig. 1F and G**). This illustrates that the rearrangement of the chromatin accessibility occurs in concert with the establishment of sex-specific gene expression programs during gonadal supporting cell differentiation.

Next, we wanted to compare our bulk RNA-seq on purified pre-granulosa and Sertoli cells (E11.5-E15.5) to a bulk RNA-seq dataset that was obtained with whole embryonic gonads (E11.5-E13.5) (*52*). It is evident that the expression levels of pre-granulosa cell markers (*e.g*. *Runx1* and *Fst)* and Sertoli cell markers (*e.g*. *Sox9* and *Amh)* differ significantly between the two datasets, showing higher expression in our data (solid lines) compared to the whole gonad data (dashed lines **Fig. 1H**) without any enrichment in other cell populations **(Fig. S4**). This is likely due to dilution of supporting cell gene expression when many other cell types are present in the whole gonad. Hence, our data constitute an important resource and the first time-series bulk transcriptomic data of the supporting cell population during the developmental window of sex determination. To facilitate further research, we offer a web application as a community tool to permit the expression level of any gene to be plotted overtime (<u>Link</u>).

These paired time-series RNA-seq and ATAC-seq data allow the in-depth characterization of the relationship between gene expression temporal dynamics and chromatin accessibility in regard to the differentiation of the pre-supporting cells into pre-granulosa and Sertoli cells.

### The sexual dimorphism of pre-granulosa and Sertoli cell transcriptomes increases with time

We undertook to characterize the transcriptomes of the differentiating supporting cells in both sexes to discern the genes that are expressed in a sex- and time-specific manner with a focus on transcription factors (TFs) as they directly contribute to gene regulatory networks.

We first conducted differential expression analysis to identify the genes exhibiting sexual dimorphism at each embryonic stage (**Fig. 2A, Fig. S5, data file S1**). Supporting cells progressively acquire a growing number of genes expressed in a sex-biased manner as they differentiate, from 1,776 and 1,424 at E11.5, to 3,713 and 3,649 at E15.5 pre-granulosa and Sertoli cells, respectively (**Fig. 2A**). The enriched GO (Gene Ontology) terms associated with sexually dimorphic genes at each embryonic stage align with our knowledge of supporting cell differentiation processes. Genes expressed at higher levels in pre-granulosa cells from E11.5 onward are involved in epithelial morphogenesis (*Tbx2*, *Bmp2*, *Bmp4* or *Shh*, to cite a few), cell differentiation and WNT signalling pathways (*Wnt2*, *Wnt11*, *Axin2* and *Snai2*, among others) (*24*, *25*, *54*), while those with higher expression in Sertoli cells from E11.5 are predominantly linked to mitotic cell cycle (*e.g. Cdk1*, *Cdc20*, *Cdc6*, *Birc5*) and epithelial morphogenesis (*e.g. Epha2*, *Sox8*, *Sox9*, *Map7*) (*55*, *56*) (**Fig. S5, data file S1**). The number of TFs exhibiting sex-biased expression also increases progressively as cells differentiate (**data file S1**). Across all stages, pre-granulosa cells exhibit higher levels of transcripts for 552 TFs compared to Sertoli cells, of these, 84 have been associated with gonadal or infertility phenotypes in the Mouse Genome Informatics (MGI) phenotype database (*57*) (**data file S1 and S2**). These include well-known critical gonadal factors such as *Tcf21* (also known as *Pod1*), *Foxl2*, *Nr0b1* (also known as *Dax1*), but also less described factors like *Osr1* that causes genital ridge hypoplasia when mutated in mice (*58*), and *Pbx3* which leads to the absence of ovaries in adults when deleted, as reported by the International Mouse Phenotyping Consortium (IMPC) (*59*). Similarly, Sertoli cells express higher levels of 471 TFs compared to pre-granulosa cells, with 76 of these associated with a gonadal phenotype upon mutation. These include well-known testicular factors *Sry, Sox9 and Dmrt1*, as well as lesser-known factors mainly associated with infertility, such as *Patz1* (*60*), *Pick1*(*61*) and *Hmga1* (*62*) (**data file S1**).

**Figure 2.**
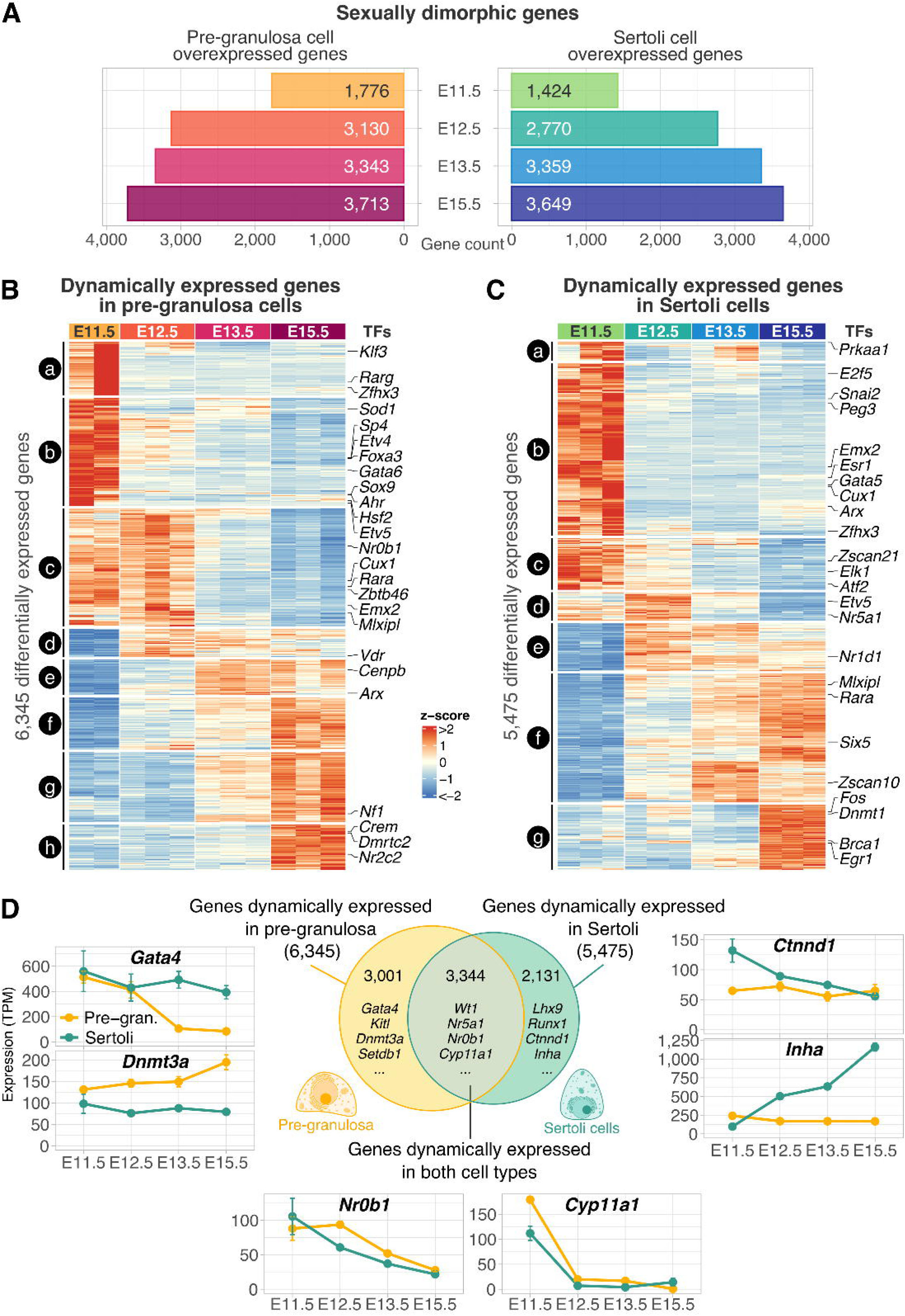
Characterization of the supporting cell transcriptome during their differentiation. (**A**) Number of overexpressed genes in pre-granulosa and Sertoli cells at each embryonic stage compared to the opposite sex. (**B**) and (**C**) Heatmaps representing the expression changes (z-score) of the genes dynamically expressed during pre-granulosa and Sertoli cells, respectively. Genes were clustered by expression profiles. The clusters were labelled with letters on the left side of the heatmaps. 25 transcription factors known to cause a gonadal or infertility phenotype were labelled on the right side of the heatmaps. (**D**) Venn diagram showing the overlap of the dynamically expressed genes in both sexes with examples of gene names present in each intersection. Expression (TPM) profiles of genes from the different sets are shown as examples.

We next explored the dynamics of the supporting cell transcriptomes in each sex separately to identify the genes whose expression changes throughout the differentiation process. Pre-granulosa cell differentiation involves 6,345 genes with expression level changes, while Sertoli cell differentiation involves modulation of 5,475 genes. These genes were subsequently classified based on their expression profiles (groups *a* to *h* in pre-granulosa cells, and *a* to *g* in Sertoli cells) and GO term enrichment analysis was performed on each of the gene profiles (**Fig. 2B and C, data file S3**). Among these dynamically expressed genes, we identified 574 TFs in pre-granulosa cells, with 89 of them reported causing a gonadal phenotype when mutated. In Sertoli cells, 530 dynamically expressed TFs were found, 86 of which are associated with gonadal phenotypes upon mutation (**data file S3**), some of them are highlighted in **Figure 2B and C**.

When overlapping the genes exhibiting expression changes during cell differentiation of both sexes, we observed that approximately half of the dynamically expressed genes in pre-granulosa cells were also found to change in Sertoli cells, and *vice versa* (**Fig. 2D**, **data file S3**). As examples, the dosage sensitive sex-determining factor *Nr0b1* (also known as *Dax1*) (*63*), and *Cyp11a1*, a cholesterol cleavage enzyme (*64*) expressed in pre-supporting cells around E11.5 (*10*) are similarly expressed in both cell types along their differentiation (**Fig. 2D**). In contrast, *Gata4* (*65*, *66*) and *Dnmt3a* (*67*) expression is relatively stable in Sertoli cells but changes in pre-granulosa cells (**Fig. 2D**). Conversely, the genes *Ctnnd1* (also known as δ-catenin) and *Inha* are stably expressed in pre-granulosa cells but change in Sertoli cells (**Fig. 2D**).

This characterization of the transcriptome during supporting cell differentiation identified a total of 802 TFs with differential expression either by sex or at specific embryonic stages as cells differentiate into pre-granulosa or Sertoli cells. Mutations of 118 of these are known to lead to a gonadal phenotype or infertility when mutated in mice, but many others have not yet been studied in the context of sex determination and gonadal development, though they may play crucial roles.

### Profiling sexually dimorphic chromatin region accessibility along supporting cell differentiation

To identify the putative *cis*-regulatory elements involved in supporting cell differentiation such as potential promoters, enhancers, silencers, and insulators, we analysed the time-series ATAC-seq data using a quantitative approach. We quantified the ATAC-seq signal of the 87,988 gonadal open chromatin regions found in all sexes and stages and assessed their level of accessibility across sexes and embryonic stages. We then performed differential accessible region (DAR) analysis to identify sexually dimorphic open chromatin regions during differentiation.

As shown in **Figure 3A**, pre-granulosa cells present fewer sex-biased accessible regions than Sertoli cells (21,608 and 31,081 in total, respectively), which is consistent with previous observations (*41*). Like the transcriptomes, the open chromatin landscape exhibits increasing sexual dimorphism as cells differentiate, from 4,206 to 16,619 regions more accessible in pre-granulosa cells and 12,679 to 24,703 regions more accessible in Sertoli cells between E11.5 and E15.5 (**Fig. 3A, data file S4**). A major proportion of the pre-granulosa-biased accessible regions (7,476) appears from E12.5 onward (**Fig. 3B**), while 9,039 Sertoli-biased regions are already established from E11.5 (**Fig. 3C**), suggesting the sex-specific accessible chromatin landscape mediating pre-granulosa cell differentiation is delayed compared to Sertoli cells, as suggested by previous transcriptomic studies (*10*, *26*). The sexually dimorphic open chromatin regions are highly enriched in intronic and intergenic regions when compared to all the open chromatin regions (**Fig. 3D, Fig. S3E**). Therefore, the sex differences in chromatin accessibility between the differentiating supporting cells are explained by the increase in accessibility of sex-specific enhancers, silencers, or insulators rather than gene promoters.

**Figure 3.**
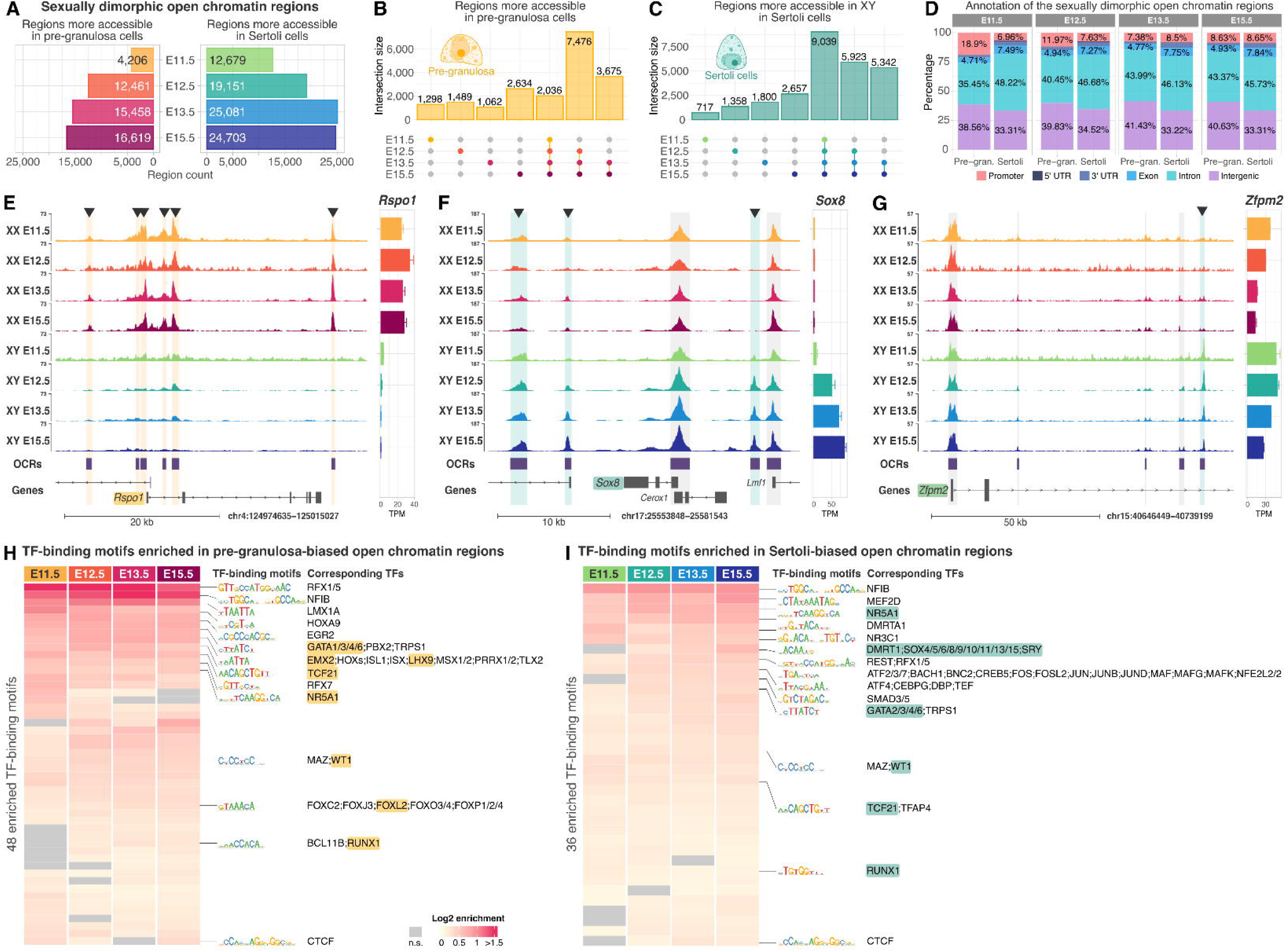
Establishment of sexually dimorphic open chromatin regions in supporting cells during their differentiation. (**A**) Number of differentially accessible regions in pre-granulosa and Sertoli cells at each embryonic stage compared to the opposite sex. (**B**) and (**C**) Upset plot representing the distribution of the sex-biased open chromatin regions across embryonic stages in pre-granulosa and Sertoli cells, respectively. Each column represents a specific intersection between sets. Dots connected by lines (bottom) indicate which sets are involved in the intersection. The height of the bars on the y-axis (top) quantifies the number of elements in each intersection. (**D**) Genomic features overlapped by the sex-biased open chromatin regions across embryonic stages in pre-granulosa and Sertoli cells. (**E**) to (**G**) Genomic tracks showing the normalized ATAC-seq signal of sexually dimorphic open chromatin regions (OCRs) around the pre-granulosa factor Rspo1, the Sertoli factor Sox8 and the critical gonadal factor Zfmp2 (also known as Fog2). Non-sex specific OCRs are highlighted in grey, sex-biased OCRs are marked by an arrowhead on top, pre-granulosa-biased OCRs are highlighted in yellow, and Sertoli-biased in blue. The bar plot on the right-hand side shows the expression level in TPM of the gene of interest. The error bars represent the standard deviation between the replicates. (**H**) and (**I**) Transcription factor binding motif enrichment analysis in pre-granulosa and Sertoli-biased open chromatin regions respectively. Only transcription factors found expressed in the supporting cells are shown. Motifs were merged by sequence similarity and the consensus logo is shown. Known gonadal transcription factors are highlighted in yellow and blue. Non-significant enrichments are coloured in grey. n.s.: non-significant.

**Figures 3E-G** exhibit examples of interesting accessible regions presenting a sex-specific pattern. The vicinity of the female-expressed gene *Rspo1* presents six genomic regions that are only accessible in pre-granulosa cells, but not in Sertoli cells (highlighted in yellow). These regions are likely to be redundantly involved in the establishment of *Rspo1* expression in pre-granulosa cells (**Fig 3E**). Likewise, exploring the *Sox8* genomic locus, we identified three male-specific regions (highlighted in light blue) which concord with the Sertoli-specific *Sox8* expression, although the *Sox8* gene promoter remains accessible in both sexes. Interestingly, our data can also identify sex-specific regions within genes expressed in both sexes. The *Zfpm2* gene (also known as *Fog2*), which is a critical co-factor of the GATA4 protein (*68*, *69*) is expressed in both Sertoli and pre-granulosa cells. While we can identify several open chromatin regions that behave similarly in both sexes (highlighted in grey), we can also identify a male-specific region located in the second intron (highlighted in light blue) (Fig. 3G). This may suggest that a gene expressed in both Sertoli and pre-granulosa cells can be regulated by different, sex-specific sets of regulatory regions.

We next looked at the enrichment of TF-binding motifs in the sexually dimorphic accessible chromatin regions. We focused our analysis on TFs that were found expressed in the supporting cells based on our RNA-seq data (**Fig. 3H and I, data file S5**). Both pre-granulosa and Sertoli-biased open chromatin regions are enriched for the motifs recognized by the non-sex-specific gonadal factors NR5A1, GATAs and WT1. Among the pre-granulosa biased open chromatin enriched motifs, we also find binding sites for known ovarian factors such as FOXL2, RUNX1, TCFs and HES1 (*70*). Similarly, we find enrichment of sites for the testis-specific factors DMRTs and SOX-SRY in the Sertoli-biased open chromatin regions. We notice, however, that the sites for GATA factors are much more enriched in pre-granulosa biased regions compared to Sertoli, and that the motif recognized by homeobox proteins, EMX2 and LHX9 which are critical factors for genital ridge development, are among the topmost enriched motifs in pre-granulosa cells (*44*, *45*, *71*, *72*).

Taken together, the results demonstrate that supporting cells undergo major sex-specific chromatin rearrangements during their differentiation. These are concomitant with the increase in accessibility of TF-binding motifs related to their respective sex-specific factors, but also factors with yet no identified role in the context of gonadal development. It remains to be deciphered whether the sex-specific chromatin accessibility changes are a cause or consequence of sex-specific TF-binding.

### Chromatin accessibility landscapes transition along supporting cell differentiation

We then focused on the open chromatin regions that change in accessibility along supporting cell development in each sex. We found 8,298 regions that change in accessibility in pre-granulosa cells and 20,607 in Sertoli cells along their differentiation, demonstrating that Sertoli cells operate a massive rearrangement of their chromatin landscape (**Fig. 4A and B, data file S6**). Among them, only 2,266 are found in common in both sexes, suggesting that the change in accessibility is mostly driven by sex-specific regions (**Fig. 4C**). We classified the chromatin regions according to the dynamics of their accessibility (groups *a* to *d* for both sexes). The chromatin accessibility events in pre-granulosa cells can be divided into two main modules. The first consists of chromatin regions that decrease in accessibility from either E12.5 (group *a*) or from E13.5 (group *b*) as cells differentiate. The second module represents chromatin regions that increase in accessibility, from E13.5 (group *c*) and from E15.5 (group *d*) (**Fig. 4A**). This suggests that the pre-granulosa chromatin landscape is transitioning from committed to differentiated cells between E12.5 and E13.5. Sertoli cells present more gradual chromatin accessibility changes, with regions from groups *a* and *b* that decrease in accessibility, and groups *c* and *d* that become more accessible (**Fig. 4B**). However, unlike pre-granulosa cells, the increase in accessibility is initiating from E12.5 onward, supporting the idea that the Sertoli specific chromatin landscape is established earlier than that of pre-granulosa cells, similar to the transcriptomic profiles (*10*, *26*).

**Figure 4.**
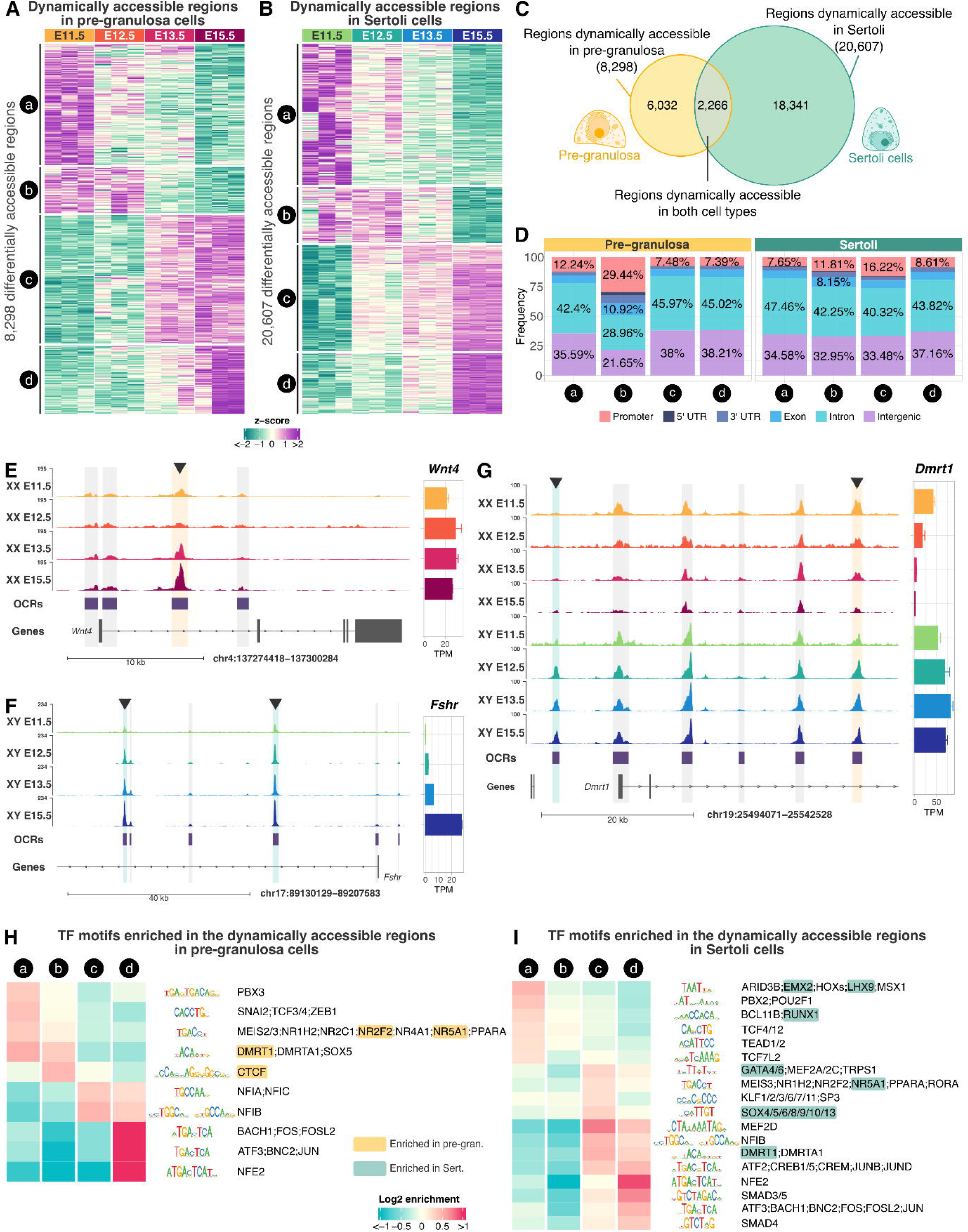
Dynamics of open chromatin regions in supporting cells during their differentiation. (**A**) and (**B**) Heatmaps representing the accessibility changes (z-score) of the open chromatin regions during pre-granulosa and Sertoli cells, respectively. Regions were clustered by accessibility profiles. The clusters were labelled with letters on the left side of the heatmaps. (**C**)Venn diagram showing the overlap between pre-granulosa and Sertoli cell dynamic open chromatin regions. (**D**) Genomic features where the dynamic open chromatin regions are found in goth pre-granulosa and Sertoli cells at each stage. (**E**) to (**G**) Genomic tracks showing the normalized ATAC-seq signal of genomic regions loci containing dynamic open chromatin regions (OCRs) in pre-granulosa and/or in Sertoli cells in the vicinity of known gonadal genes. Non-dynamic OCRs are highlighted in grey, significantly dynamic OCRs are marked by an arrowhead on top, pre-granulosa-dynamic OCRs are highlighted in yellow, and Sertoli-dynamic in blue. The bar plot on the right-hand side shows the expression level in TPM of the gene of interest. The error bars represent the standard deviation between the replicates. (**H**) and (**I**) Heatmap of the transcription factor motifs differentially enrichment between the different open chromatin region clusters from pre-granulosa and Sertoli cells, respectively. Only transcription factors found expressed in the supporting cells are shown. Motifs were merged by sequence similarity and the consensus logo is shown. Known gonadal transcription factors are highlighted.

These genomic regions are mainly located in intronic and intergenic regions as observed with the sexually dimorphic accessible regions (**Fig. 4D**). **Figure 4E-F** presents several interesting examples of changes in chromatin accessibility over time. For example, we observe that the *Wnt4* gene, which is expressed in pre-granulosa cells, presents an open chromatin region in its first intron that increases in accessibility as cells differentiate (highlighted in yellow), while the promoter of the *Wnt4* gene remains stably accessible over time (**Fig. 4E**). Similarly, we identified two intronic open chromatin regions on the *Fshr* male specific gene that increase in accessibility as cells differentiate and are likely to be putative enhancers (**Fig. 4F**). Among the dynamically accessible regions present around gonadal critical gonadal genes, we could observe more complex patterns. *Dmrt1*, which is expressed in both pre-granulosa and Sertoli at E11.5 and become exclusively expressed in Sertoli cells from E12.5 onward, presents one open chromatin region upstream of the promoter that gradually becomes more accessible specifically in Sertoli cells (highlighted in light blue), while another region in the second intron shows a decrease in accessibility in pre-granulosa cells while remain stably accessible in Sertoli cells (highlighted in yellow) (**Fig. 4G**). Altogether this suggests a complicated gene regulatory network both temporally and also between sexes.

The changes in the open chromatin landscape also reflect a change of accessibility of specific binding motifs from TFs expressed in pre-granulosa and Sertoli cells (TPM>10). Differential TF-binding motif enrichment analysis showed that pre-granulosa cell regions that are more accessible at early stages (groups *a* and *b*) are enriched for motifs recognized by NR2F2 and NR5A1, but also by DMRT1. This concords with the fact that supporting cells are derived from *Nr2f2* and *Nr5a1* expressing progenitor cells, while the expression of both of these genes decreases as cells differentiate into pre-granulosa cells (*9*, *10*), and the fact that *Dmrt1* is expressed in E11.5 pre-granulosa cells (**Fig. 4G**). Group *d* is strongly enriched for motifs recognized by different factors such as FOS (**Fig. 4H, data file 7**). In Sertoli cells (**Fig. 4I, data file 7**), we see that group *a* regions, *i.e.* those more accessible at E11.5, are enriched in motifs for LHX9 and RUNX1, TFs whose expression is down-regulated in Sertoli cells after E11.5 (*9*, *42*). Group *c* regions, which are gradually more accessible from E12.5, are enriched in motifs for GATA, NR5A1, DMRT1, and SOX proteins; and group *d* regions also show enrichment for DMRT1, a TF known to be crucial for sex maintenance in the testis.

Taken together, the ATAC-seq data analysis of the differentiating pre-granulosa and Sertoli cells allowed the identification of genomic loci presenting a difference in accessibility between sexes and embryonic stages. These regions are enriched in motifs for known critical sex-specific TFs but also many others that have not yet been described in the context of gonadal development. These data also provide a wide atlas for the identification of stage- and sex-specific regulatory elements that may be involved in the tight and precise regulation of gene expression during sex determination, mutations in which may lead to DSDs.

### Predicting the target genes of the putative supporting cell *cis*-regulatory regions

After having characterized the transcriptome and the open chromatin landscape of the supporting cells, we set out to predict the target genes of the putative *cis*-regulatory elements. One of the most commonly used proxies to achieve this is to look for the correlation between gene expression and accessibility of the open chromatin regions in the +/−500 kb region from their TSS. A positive correlation, or link, corresponds to a chromatin region accessibility that mirrors the nearby gene expression and therefore suggests a putative enhancer function. Conversely, a negative correlation means that the chromatin accessibility is the opposite of the nearby gene expression suggesting it may function as a silencer (**Fig. 5A**). Using this method, we could identify 44,116 putative regulatory regions (50.1% of all the open chromatin regions) whose accessibility correlates positively or negatively with 10,685 genes, which represents 88.6% of all the expressed protein coding genes (**data file S8**). Each gene was linked to an average of seven putative *cis*-regulatory elements, four positively and three negatively (**Fig. 5A, Fig. S6**) and each linked open chromatin region was connected to a median of two genes (**Fig. S6**). The linked open chromatin regions are mainly located within intragenic regions, including within the gene with which they are linked, and intergenic regions (**Fig. S6**).

**Figure 5.**
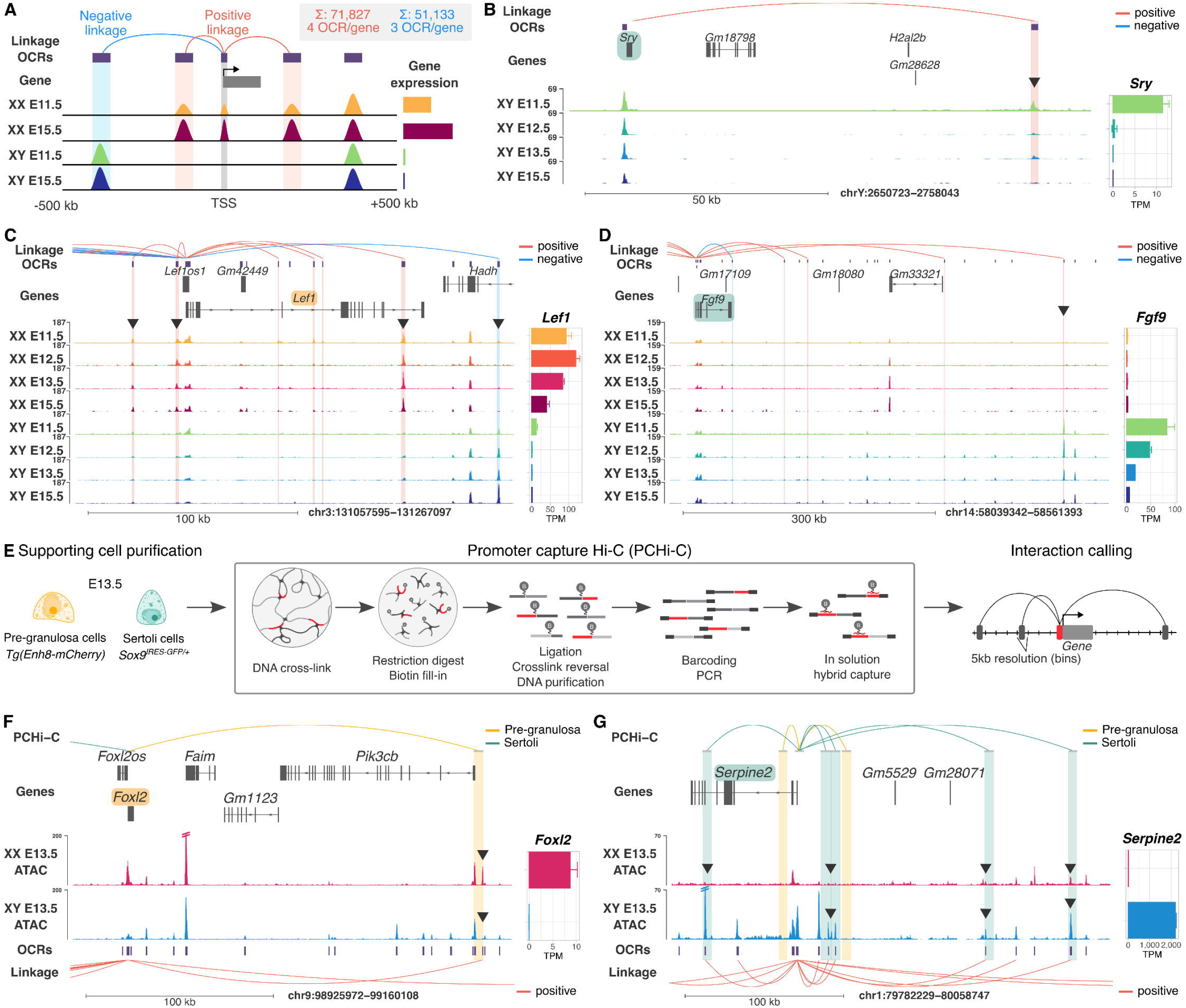
Prediction of gene-cis-regulatory region associations. (**A**) Schematic representation describing the strategy to link open chromatin regions with their target genes. Open chromatin regions located within a ±500 kb window of a gene’s transcription start site (TSS), whose accessibility correlates with gene expression, are identified as putative enhancers. Conversely, open chromatin regions that anti-correlate with gene expression are linked as putative silencers. The number of positive and negative link, as well as the average number of linked open chromatin regions (OCR) per gene are indicated. (**B-D**) Genomic tracks showing the predicted links between open chromatin regions and gene expression. The positive links are represented as red line arcs, and negative links as blue line arcs. The genomic tracks represent normalized ATAC-seq signal of genomic regions loci in pre-granulosa and/or in Sertoli cells. Noticeable genomic loci are indicated with an arrowhead. The bar plot on the right-hand side shows the expression level in TPM of the gene of interest. The error bars represent the standard deviation between the replicates. (**E**) Experimental design of the promoter capture Hi-C (PCHi-C) experiment. E13.5 pre-granulosa and Sertoli cells have been purified by FACS, fixed, and have been subjected to PCHi-C. Interactions between gene promoters and genomic regions were called using a 5 kb bin resolution. (**F-G**) Genomic tracks showing the PCHi-C interactions found in either pre-granulosa or Sertoli cells around two sexually dimorphic genes at E13.5. The interactions contain open chromatin regions and overlap with the RNA-ATAC linkage analysis. The dash lines on top of the ATAC-seq signal signify that the scale has been cropped.

In **Figures 5 B, C** and **D**, we show three examples of sex-specific genes involved in gonadal development and some of their regulatory elements that present either a positive or negative correlation. We found only one open chromatin region that was linked to the testis determining gene *Sry*, located 84 kb upstream of its TSS (**Fig. 5B**). This region corresponds to the locus containing transcriptionally active sequences derived from transposable elements we previously identified (*73*). *Lef1*, a downstream gene of the WNT signalling, presents 15 open chromatin regions that correlate with its expression in a window of +/−500 kb from its TSS (**data file S8**). These include 10 putative enhancers, *i.e.* open in pre-granulosa but not in Sertoli cells, and five putative silencers, *i.e.* close in pre-granulosa but open in Sertoli cells. **Figure 5C** shows two upstream and four intronic open chromatin regions that are likely to be *Lef1* enhancers, and one downstream region with accessibility that anti-correlates with *Lef1* expression and could act as a silencer (**Fig. 5C**). Similarly, we found nine putative *Fgf9* enhancers, and one potential silencer (**data file S8**). **Figure 5D** shows four distal downstream putative *Fgf9* enhancer regions and a potential downstream silencer (**Fig. 5D**). Interestingly, deletion of a 306 kb region that includes the most downstream enhancer resulted in XY male-to-female sex reversal in mice (*74*).

To complement the linkage prediction analysis, we performed promoter capture Hi-C (PCHi-C) on purified E13.5 Sertoli and pre-granulosa cells (**Fig. 5E**). PCHi-C enables to profile chromatin interactions between promoters of protein-coding genes and their regulatory elements (*75*, *76*). Significant contacts were detected with CHiCAGO (*77*) at a 5 kb resolution using two approaches (original bait and 5 kb extended bait) according to the best practice guidelines, with and without inclusion of promoters in the binning process to increase the detection sensitivity for proximal and distal interactions (*75*) (see Methods, **Fig. S7A**). In total, we detected 82,532 and 108,206 contacts (interaction score > 3) in pre-granulosa and Sertoli cells, respectively, between promoters and promoter-interacting regions (PIRs) at 5 kb resolution, 60,909 of them being commonly found in both sexes (interaction score > 3 in both sexes) (**data file S9**). Pre-granulosa and Sertoli interactions were enriched for markers of accessible and/or active enhancers (ATAC, H3K27ac) and active transcription (H3K4me3), previously described in pre-granulosa and Sertoli cells (*41*) (“active PIRs”, **Fig. S7B**). Moreover, we observed a positive correlation between mean gene expression and the number of promoter-interacting regions, which suggest that highly expressed genes tend to be controlled by more *cis*-regulatory regions (**Fig. S7C**).

Overall, we found 3,603 PCHi-C interactions that overlap with 3,484 links predicted with our linkage analysis. **Figure 5F and G** show examples of physical interactions that we also predicted around two sexually dimorphic genes. We identified a physical interaction between a distal pre-granulosa-specific open chromatin region downstream of *Foxl2* and its promoter (**Fig. 5F**), strongly suggesting this region functions as an enhancer for *Foxl2*. Similarly, we observed five Sertoli-specific open chromatin regions that physically interact with the *Serpine2* gene, highly expressed specifically in Sertoli cells (**Fig. 5G**).

This data provides the first promoter-targeted chromatin interactome in Sertoli and pre-granulosa cells, shedding light on the regulatory signalling involved in sex determination.

### Different sets of TFs are physically bound to regulatory elements in pre-granulosa and Sertoli cells

For deciphering comprehensive gene regulatory networks, it is crucial to identify regulatory elements, physically associate them to their target genes and identify the TFs that bind these enhancers. While we identified the accessible chromatin regions in pre-granulosa and Sertoli cells and were able to predictively and physically associate them to their target genes, we still aim to identify which TFs bind to these regulatory regions to control precise gene expression patterns. While few TF ChIP-seq experiments were performed on gonadal cells (*28*, *49*, *50*, *78*), it remains a major challenge due to the highly limited number of gonadal cells at embryonic stages. To overcome this challenge and identify potential TFs that are physically bound to accessible chromatin regions, we conducted ATAC footprinting analysis on the open chromatin regions identified in both cell types at all developmental stages.

To that aim, we first looked at the enrichment of TF-binding motifs in the sexually dimorphic accessible chromatin regions. We performed differential TF-binding motifs enrichment on the sexually dimorphic accessible chromatin regions to detect motifs that are more accessible in one sex compared to the other (**Fig. 6A, Fig. S8, data file S5**). Next, we used the ATAC-seq data to detect TF-binding motif occupancy or footprinting. This represents physical binding of protein onto the open chromatin DNA in a way that confers small blockage to Tn5 digestion (*79*). We measured the difference of occupancy of the motifs of the expressed TFs between pre-granulosa and Sertoli cells in the sex-biased open chromatin regions and confirmed that most of the sexually dimorphic enriched motifs are also differentially occupied by a transcription factor (**Fig. 6A, data file S10**). Surprisingly, although the RFX1/5 and ZBTB14 motifs are among the top three enriched transcription factor motifs of pre-granulosa cell-biased open chromatin regions compared to Sertoli cells, they are not the most differentially bound motifs. The motif recognized by EMX2, LHX9 and MSX1, among others, is the most differentially bound compared to Sertoli (**Fig. 6A and B**), and its score increases with time, suggesting that these factors are potentially the most important for the establishment of the pre-granulosa specific gene regulatory network. Conversely, the motifs recognized by the DMRT1 and SOX TFs are the most differentially bound in Sertoli cells (**Fig. 6A and C**), confirming their key role in controlling Sertoli cell differentiation and maintenance (*3*, *80*).

**Figure 6.**
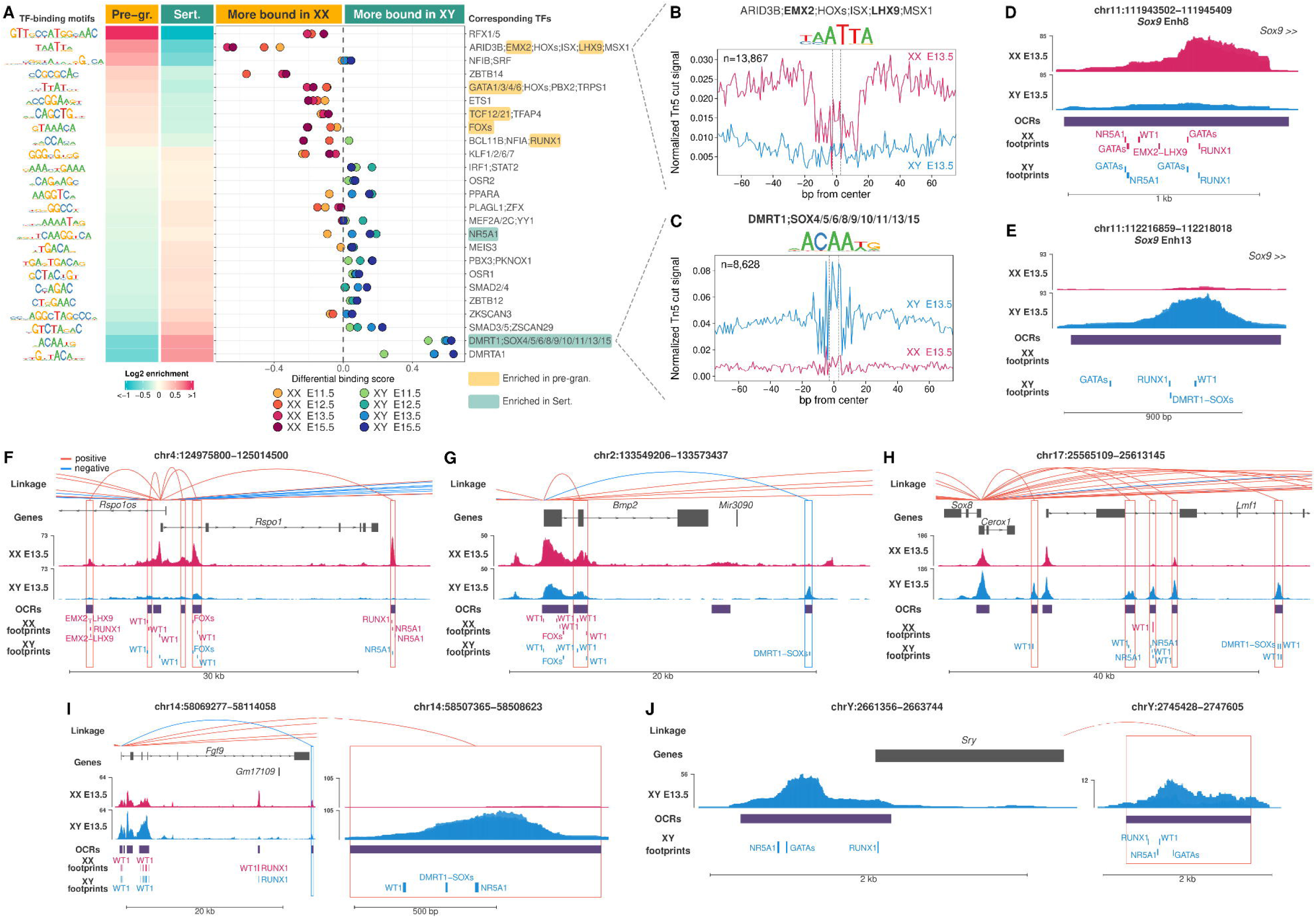
ATAC TF footprints on sexually dimorphic open chromatin regions. (**A**) Differential transcription factor binding motif enrichment and occupancy (ATAC-seq footprints) in the combined (all developmental stages) sex-biased open chromatin regions when comparing one sex to the other. Only transcription factors found expressed in the supporting cells are shown. Motifs were merged by sequence similarity and the consensus logo is shown. Known gonadal transcription factors are highlighted. (**B**) and (C) Comparison of the aggregated ATAC-seq footprint signals at E13.5 in both sexes for the top pre-granulosa and Sertoli cell differentially bound TF-binding motifs recognized by EMX2 and LHX9, and DMRT1 and SOX and SRY factors, respectively. The number of bound motifs is indicated. The dashed lines represent the motif location. (**D**) to (**J**) Genomic tracks showing TF footprints of the gonadal factors highlighted in figure (**A**) and WT1 on different sex-biased accessible loci. Footprints found in any of the studied embryonic stages were aggregated. For concision, we only show the name of the known gonadal TFs and not the full list of TFs able to bind the occupied motifs.

We focused our interest on the factors that are the most differentially bound in the pre-granulosa cell-biased open chromatin regions (ARID3B, EMX2, LHX9, ISX and MSX1). We checked the expression level of their genes in our RNA-seq data (**Figure S9A**) and in the single-cell RNA-seq atlas of the developing gonad (*9*) (**Figure S9B**) to identify which factors are the most highly and specifically expressed in the pre-granulosa cells. We found that *Arid3b* and *Isx* are lowly expressed in pre-granulosa cells in both datasets (**Figure S9**). *Msx1* is detected at a higher level in our bulk RNA-seq (**Figure S9A**) than in the single-cell data (**Figure S9B**), and is also expressed in the female germ cells (*9*). Finally, *Emx2* and *Lhx9*, while they are expressed in both sexes in the early progenitors (prior to supporting cell commitment) and the pre-supporting cells (E11.5 supporting cells), their expression decreases in Sertoli cells and become almost restricted to pre-granulosa cells (**Figure S9**). These observations suggest that EMX2 and LHX9 could play a crucial role in the pre-granulosa cell commitment.

Finally, we looked at TF footprints at a locus resolution around known enhancers like Enh8 and Enh13 of the *Sox9* gene (**Fig. 6D and E**), but also within candidate enhancers of sex determination genes to identify potential gonadal factor binding sites (highlighted in **Fig. 6A**) (**Figure 6D** to **J**, **data file S10**). We can see footprints likely to be associated with binding of SOX factors to Enh13 (**Fig. 6D**) as well as of several pro-female factors to Enh8 as RUNX1, GATAs, NR5A1 and LHX9/EMX2 (**Fig. 6E**). We can identify footprints of WT1 on open chromatin region in *Rspo1* promoter and first intron (**Fig. 6F**) that overlap with WT1 ChIP-seq data performed on embryonic kidneys. This may suggest that shared regulatory elements are exploit in the kidney and gonads and that *Rspo1* is a direct target of WT1 (*81*). We also see WT1, SOX/DMRT and NR5A1 footprints to the downstream enhancer of *Fgf9* gene (**Fig. 6I**) but also footprinting of WT1, GATAs, NR5A1 and RUNX1 footprints on the upstream enhancer of *Sry* (**Fig 6J**). This analysis can pinpoint the key factors likely to bind each regulatory element of each gene.

Altogether, we find that the sex-biased open chromatin regions are enriched in TF motifs for the known sex determining factors of the respective sex, but also for other TFs that had not previously been identified as having a role in gonad development. Thus, these regions constitute putative genomic targets of the sex-determining factors. Furthermore, it is possible that factors with known function at early gonad development such as EMX2 and LHX9 also play a critical role at later stages during pre-granulosa cell development.

## Discussion

Decades of research on mouse sex determination and human DSD individuals have unravelled many of the TFs and signalling pathways involved in the process. Yet, it is striking that ∼70% of the 46,XX DSD and ∼50% of 46,XY DSD patients fail to receive genetic diagnosis following whole exome sequencing (*82*, *83*). This suggests that variants in the 98% of the non-coding genome are highly likely to explain many of the unresolved DSD cases. This emphasises the need to identify the *cis*-regulatory elements that control the antagonistic gene networks at play during mammalian sex determination.

In this study, our aim was to decipher the *cis*-regulatory elements that participate in the process of gonadal supporting cell differentiation, the genes they regulate, and the transcriptomic outcome. To achieve this, we purified fetal pre-granulosa and Sertoli cells using the newly generated Enh8-mCherry and the established *Sox9^IRES-GFP^* mouse lines at four embryonic stages covering the process of sex determination and subsequent supporting cell differentiation. One limitation of the use of *Sox9^IRES-GFP^*to purify Sertoli cells is that we preferentially enriched our multiomics data with already committed Sertoli cells at E11.5, as shown by the low expression of *Sry* in our dataset compared to whole gonads (**Fig. S4C**). However, the use Enh8-mCherry and the *Sox9^IRES-GFP^* mouse lines allowed us to increase our cell sorting yield by five-fold compared to the previously used *TESCO*-CFP and *TESMS*-CFP transgenic mouse lines (*40*, *41*) (**Fig. S2D**). Moreover, the proportion of cells we obtained using these two mouse lines is close to what was observed in the whole gonads, according to scRNA-seq data (*9*). This suggests that our study is based on representative fetal gonad supporting cell populations.

In recent years, several studies have generated time-course transcriptomic profiles of mouse gonadal development using both whole gonad bulk RNA-seq (*49*, *52*, *84*) and scRNA-seq (*9*, *10*, *42*). While these studies have been transformative to the field and enabled the identification of novel and rare cell populations, they also hold some limitations. Bulk RNA-seq of whole gonads lacks the resolution needed to characterize gene expression within specific gonadal cells, a challenge that can be overcome by scRNA-seq technologies. However, scRNA-seq techniques are limited in sensitivity, making it difficult to detect low-abundant transcripts (*85*). Our bulk RNA-seq data represent the first transcriptomic analysis of purified Sertoli and pre-granulosa cells during early gonad development. Although averaging cells in which differentiation is asynchronous, the sensitivity of bulk RNA-seq makes this method as a good complement to scRNA-seq data for lowly expressed genes.

Previous ATAC-seq analysis was performed on E10.5 supporting cell precursors and E13.5 Sertoli and pre-granulosa cells (*41*) and E14.5 and later pre-granulosa cells (*49*), leaving a big gap of the most critical stages in which gonadal sex is being established. Here, we performed RNA-seq, as well as ATAC-seq on four critical developmental stages covering the entire window of sex determination. Overall, we found that 2.8% of the 98% non-coding DNA are accessible during male and female supporting cell differentiation. Our analysis indicates that while many *cis*-regulatory elements exhibit sexual dimorphism in accessibility staring at E11.5 in Sertoli cells, most pre-granulosa-biased open chromatin regions are established from E12.5, reinforcing the idea that pre-granulosa cell commitment is delayed compared to that of Sertoli cells (*10*, *32*, *47*, *86*).

TF-binding motif enrichment analysis and footprinting indicate that most of the Sertoli-cell biased regions are bound by DMRT and SOX factors. SOX9 has been described as being a pioneer TF that is able to bind closed chromatin, inducing *cis*-regulatory element opening that leads to cell reprogramming in embryonic epidermal stem cells and umbilical vein endothelial cells (*87*, *88*). All these data confirm the importance of SOX9 as the master regulator for the onset of Sertoli cell fate decision by establishing the chromatin landscape necessary for the activation of the male-specific gene program. It was also shown that ectopic DMRT1 in the ovary acts as a pioneer factor to induce *Sox9* expression but also repress *Foxl2* (*89*, *90*). Interestingly, depletion of *Dmrt1* in males does not affect testicular development (*91*), which argues in favour of a role in the maintenance of Sertoli cell identity rather than a determining gene. However, its expression in E11.5 pre-granulosa cells and the enrichment of its binding motif in early pre-granulosa cells (when *Foxl2* is not yet expressed), can fit with its role to negatively regulate *Foxl2,* and, more globally, female-specific gene expression. This data may also potentially explain the delay in pre-granulosa cell differentiation compared to Sertoli cells.

Regarding pre-granulosa cells, we observed that RUNX1 and FOX motifs were not the most enriched in the pre-granulosa-biased open chromatin regions at the onset of their differentiation. This is consistent with the fact that deletion of these TFs in fetal gonads leads to a delayed female-to-male sex reversal, either in postnatal ovaries when only *Foxl2* is deleted (*31*), or in late fetal stages, around E15.5, when both factors are deleted (*28*). As such, FOXL2 and RUNX1 are required for granulosa cell fate maintenance rather than being ovary determining factors. However, our study sheds light on two factors that may act on pre-granulosa cell commitment: EMX2 and LHX9. While they have been shown to be crucial for bipotential gonad formation (*44*, *45*, *92*), their role at later stages has never been characterized. The identification of the *Wt1^-KTS^* isoform as the female determining factor is a clear illustration that early factors can also have critical roles at later stages (*26*). *Emx2* and *Lhx9* expression become restricted to the pre-granulosa cells soon after the supporting cell commitment, from E11.5 (*9*, *10*, *47*, *93*, *94*). Concomitantly, the pre-granulosa cell accessible genome presents increasing enrichment and binding footprints of the motifs recognized by these two factors. Although our analysis cannot decipher whether one or both are actually binding on the pre-granulosa putative *cis*-regulatory elements, EMX2 and LHX9 represent the most promising candidate TFs that could regulate the pre-granulosa cell differentiation genetic program. Moreover, EMX2 has been shown to play an important role in mouse neuroblast proliferation, migration and differentiation (*71*), while LHX9 has not been identified in cell fate decision role in other systems.

Integration of both RNA and ATAC-seq data allowed for the linking of putative *cis*-regulatory elements to their target genes, allowing the identification of thousands of pre-granulosa- and Sertoli-specific putative *cis*-regulatory elements. Yet, it is likely that this is an understatement of the real state, as we limited the linkage analysis to +/− 500 kb, and critical enhancers like Enh13 (*Sox9*) and ZRS (*Shh*) are often located over 500 kb away from their target genes (*37*, *95*). Similar findings in other systems suggest a complex regulatory network of enhancers and silencers, many acting redundantly to control gene expression. While redundancy is common, single enhancers like Enh13 and ZRS can fully regulate target genes at specific stages, with deletions causing dramatic phenotypes.

Our PCHi-C analysis corroborated some of the *cis*-regulatory element-gene association predicted by the linkage analysis. Yet, while able to detect specific enhancer-promoter interactions, we could not detect the interaction of Enh13 to the *Sox9* promoter in Sertoli cells. This suggests that the PCHi-C, while revealing interesting interactions, is not comprehensive enough and needs to be further refined. Indeed, PCHi-C is normally being performed using millions of cells, and here it was done using 100K sorted gonadal cells. It is possible that using a larger number of cells could improve the resolution.

Altogether, this study serves to unravel the gene regulatory networks that are at play during mammalian sex determination. It uncovers the *cis*-regulatory elements that function during sex determination, along with the binding motifs enriched and occupied in them, the physical binding of regulatory elements and target genes, and how all this eventually translates into unique gene expression profiles. To complete our understanding of the gene regulatory networks, it will be important to have additional layers of information such as histone post-transcriptional modifications as well as binding of TFs and histone modifiers.

## Materials and Methods

### Mice and ethics

All animals were maintained with appropriate husbandry according to Bar-Ilan University ethics protocols 11-02-2020, 57-08-19, 2305-111-1 and 2306-117-1. All mouse strains were maintained on an F1 (C57BL6/J x CBA-Ca) genetic background. *Tg(Enh8-hsp68-mCherry* mice were generated at the Francis Crick Institute transgenic facility by zygote microinjection and imported to Israel.

The X-GFP (*Tg*(*CAG-EGFP*)*D4Nagy*) (*96*) and *Sox9^IRES-GFP/+^* (*53*) mice were used. Embryos and animals used in this study were either *Tg(Enh8-hsp68-mCherry)* heterozygotes for Cherry or *Sox9^IRES-GFP^* heterozygotes for GFP. Primers used for genotyping these mice strains are listed in **Table S1**.

### Cloning of Enh8-hsp68-mCherry vector

Enh8 (mm9: 111,805,552-111,806,224; 672 bp long, located 838 kb upstream of the *Sox9* gene) was amplified by PCR and cloned into the AseI restriction sites of the pmCherry-N1 vector (Kan resistance) reporter vector (Takara Cat No. 632523) using In-Fusion HD (Clontech). The *hsp68* sequence was amplified from the pSfi-Hsp68-LacZ reporter vector (Addgene #33351) via In-Fusion HD and cloned into the AseI and AgeI restriction sites of the pmCherry-N1 vector. The primers used for the cloning are presented in **Table S1**. To release the entire Enh8-hsp68-mCherry construct, the AseI and NotI restriction enzymes were used. All plasmids were verified using Sanger sequencing.

### Generation of the *Tg(Enh8-mCherry)* mice

To prepare DNA for zygote injection, 50 μg of the Enh8-hsp68-mCherry plasmid was digested with AseI and NotI and the insert was gel purified by electroelution. The DNA was phenol-chloroform extracted, ethanol precipitated and resuspended in TE buffer (10 mM Tris-HCl, 1 mM disodium EDTA, pH 8.0). The DNA was further purified on a DNA-cleanup column (Qiagen PCR Purification Kit). 5 ng/μl of the purified plasmid was injected into pronuclei of F1 (C57BL/6JxCBA) zygotes. These were transferred on the same day into the oviducts of pseudopregnant CD1 females. Injections were performed by the Crick Genetic Manipulation Service to generate stable transgenic lines. Cherry positive founders were bred to wild type (C57BL/6JxCBA) F1 females to first examine expression profiles in the gonads of E13.5 embryos and then perform germline transmission. *Tg(Enh8-hsp68-mCherry)* line No. 3 was found to show robust expression in the gonads and was bred for a few generations to “clean” the line from extra transgenes until a stable line was established. This line is termed Enh8-mCherry (*Tg(Enh8-hsp68-mCherry))*.

### gDNA isolation and transgenic mice genotyping

Genomic DNA (gDNA) was extracted from tail tissue of embryos or ear punch tissue of adult animals. gDNA isolation from adult earpiece included 15 min incubation at 95°C with lysis buffer composed of 10 mM NaOH, 0.1 mM EDTA pH 8 followed by the addition of 40 mM Tris-HCl pH 5. For embryo samples, the PCRBIO rapid extract lysis kit was used (PCRBIO, PB15.11-S).

All PCR reactions were performed using 2X of Dream-Taq PCR mix (Thermo, K1082) according to the manufacturer’s instructions. All mice and embryos were also genotyped for the chromosomal sex (*97*) (Genotyping primers are listed in **Table S1**).

### Time mating, gonad harvesting, tissue preparation and imaging

Embryos and animals carrying the desired transgene were produced by crossing a *Sox9^IRES-GFP^* male or a *Tg*(*CAG-EGFP*)*D4Nagy; Tg(Enh8-hsp68-mCherry)* male mice on F1 (C57BL/6J x CBA) genetic background female mice. Pregnant females were sacrificed by CO_2_ inhalation and embryos were harvested at different stages of the pregnancy (E11.5, E12.5, E13.5 and E15.5). Day 0.5 was determined by the presence of a vaginal plug (VP). For E11.5 or E12.5 gonads, we considered the embryos that had between 18 and 21 tail somites (TS) as E11.5 and embryos that had 27-30 TS as E12.5. The sex of gonads at E13.5 and E15.5 was determined via visual inspection. The sex of the E11.5 and E12.5 *Tg*(*CAG-EGFP*)*D4Nagy; Tg(Enh8-hsp68-mCherry)* embryos (where the Cherry appears in both XX and XY) was determined by having the X-GFP transgene where GFP-positive embryos are XX and GFP-negative embryos are XY (*96*).

All bright field and fluorescent images of gonads were taken using the Nikon Eclipse Ts2R microscope. Exposure times: 10 ms for BF, 5-7s for mCherry, 2-3 seconds for GFP. Images were analysed using the NIS-Elements D software.

For immunostaining, gonads from embryos were harvested and fixed overnight in 4% paraformaldehyde (PFA) (Sigma Aldrich, P6148) in phosphate buffered saline (PBS) at 4°C, washed three times with PBST (PBS with 0.1% Triton (Sigma Aldrich, 9002-93-1)) at room temperature, incubated with 20% sucrose (Fisher BioReagents, BP220-1) overnight at 4°C and then embedded in OCT (Leica, 14020108926) and stored at −80°C until further use.

### Immunofluorescence staining

Immunofluorescence staining was performed on 10 μm-thick sagittal cryostat sections (Leica, CM3050-S). Antigen retrieval was performed with DAKO (Target retrieval solution, Agilent, S1699) at 65°C for 30 min. Samples were then blocked in PBST containing 10% donkey serum (Sigma Aldrich, D9663) for 1hr and incubated with primary antibodies (diluted in PBST containing 1% donkey serum) overnight at 4°C (All primary and secondary antibodies as well as dyes used are listed in **Table S2**). Following three PBST washes, secondary antibodies were added and incubated for 1hr at room temperature (RT). Slides were then washed, dried and mounted (Polysciences 18606-20). All immunofluorescence slides were also stained with 4, 6-diamidino-2-phenylindole (DAPI; Invitrogen, D1306) to visualize nuclear DNA. Images were obtained with a Leica Microsystems SP8 confocal microscope.

### FACS of Sertoli and pre-granulosa cells

Following dissection, gonads were separated from the mesonephros using a G25 needle and dissociated into single cells by culture in 0.045% trypsin-EDTA (Thermo, cat. 25300062) along with 0.25% collagenase (Wortington, cat. LS004176) in a total volume of 500 µl for 8 min at 37°C in a 24-well plate. 200 µl of DPBS (without Calcium and Magnesium) containing 3% Bovine serum albumin (BSA) (Sigma-Aldrich, cat. A3311) were added to quench the trypsin action and immediately aspirated to remove the trypsin. Gonads were then resuspended in 300µl of DPBS containing 3% BSA and mechanically dissociated by gentle pipetting. Cells were filtered through a 30µm strainer (Miltenyi Biotec, cat. 130-041-407) and collected into a Polypropylene FACS tube (BD Falcon, cat. 352063). The well and filter are washed with additional 200µl of DPBS containing 3% BSA. Cells were sorted using the BD FACS Aria III cell sorter with an 85-micron nozzle. Texas Red filter (excitation 561 nm, emission 610 nm) was used to detect mCherry and FITC (excitation 495 nm, emission 519 nm) filters to detect GFP. For ATAC sequencing, positive cells were sorted into Eppendorf tubes containing 150µl of DPBS + 3% BSA. For RNA sequencing positive cells were collected into 150µl QIAzol lysis reagent (Qiagen, 79306). FACS raw data were analysed using FlowJo v10 for visualization (**Fig. S2A-C**).

### RNA-seq and library prep

Total RNA was isolated from ∼40,000 sorted Sertoli or pre-granulosa cells of E11.5, E12.5, E13.5 and E15.5 gonads via FACS. RNA purification was performed using the RNeasy plus micro kit (Qiagen, 74034) according to the manufacturer instructions. cDNA libraries were prepared using the NEB Next Single Cell/Low Input RNA Library Prep Kit for Illumina (NEB, E6420) according to the manufacturer’s protocol. cDNAs were synthesized by reverse transcriptase enzyme, followed by cDNA amplification and cDNA clean-up using AMPure beads (Beckman coulter, Bc-a63881). cDNA was then fragmented, and adapters were ligated with unique indexes for each sample using the NEBNext® Multiplex Oligos for Illumina (NEB, E7335). cDNA and library concentrations were analysed using Qubit ds HS Assay Kit (Invitrogen, 2326054) and sample size distribution using Tapestation with high sensitivity D1000 tape (Agilent, 5067-5585). Samples were pooled together to create a 4nM library and sequenced with 83 bp single end (SE) reads on the Illumina Next-Seq 500 platform at the Bar-Ilan Sequencing Unit with roughly 25-30 million reads per sample.

### RNA-seq mapping and quality controls

FastQ files were processed using nf-core/rnaseq pipeline v3.12.0 (*98*). Briefly, read quality controls were performed with FastQC. Sequencing adapters were removed with TrimGalore!. Reads were mapped on the mm10/GRCm38 reference genome from Gencode (M25) with STAR. Gene quantification as read counts and TPM (Transcript Per Million) was obtained using RSEM. Read count matrix was filtered to exclude lowly expressed genes (genes with less than 15 reads and/or TPM value less than 5). Non-protein-coding genes were filtered out in the subsequent analysis. Sample correlation (Spearman) and PCA were performed with R (corr and prcomp functions) using the filtered read counts normalized by library size (sizeFactor) with DESeq2 (*99*). After inspection of the correlation and the PCA, we excluded the XX E11.5 replicate 2 because it was more similar to XX E12.5 samples than the E11.5 samples. This is probably due to embryos that were older than expected the day of the collection (**Fig. S3A-B**).

### Differential expression analysis

Sex differential expression analysis was performed on the filtered read count matrix using DESeq2 with ∼Sex_stage as model and using the “Wald” test. The number of sexually dimorphic genes per stage was retrieved using the contrasts. Genes with log2(FoldChange) < −0.5 and log2(FoldChange) > 0.5, as well as an adjusted *p*-value < 0.01 were considered as differentially expressed.

Stage differential expression analysis was performed on each sex separately using DESeq2 with ∼Stage as model and using “LRT” (Likelihood Ratio Test). As previously, genes with log2(FoldChange) < −0.5 and log2(FoldChange) > 0.5, as well as an adjusted *p*-value < 0.01 were considered as differentially accessible. The overlap of the sex differentially expressed genes across stages was computed and represented as an upset plot using eulerr and UpsetR packages (*100*, *101*).

Gene expression values (filtered read count matrix normalized by library size) from the stage differentially expressed genes were transformed as z-scores, clustered into groups according to their expression profiles using hclust and “ward.D2” method, and represented as heatmap using ComplexeHeatmap (*34*). The optimal number of clusters were assessed using the best.cutree function from the Jlutils R package (https://github.com/larmarange/JLutils).

The overlap between the sex and the stage differentially expressed genes was computed and represented as a venn diagram using the eulerr package. The gene expression visualisation website was developed using Shiny and Plotly R packages.

### GO term enrichment analysis

Gene Ontology enrichment analyses were performed using ClusterProfiler (*102*). Biological process GO terms with an enrichment *p*-value < 0.01 and *q*-value < 0.05 were reduced by similarity using the “simplify” function with a similarity cutoff of 0.7.

### TF and phenotype annotation

Differentially expressed genes were annotated whether they are transcription factors and whether a mutated allele in mice induced a gonadal or fertility-related phenotype.

The list of mouse transcription factors was taken from https://resources.aertslab.org/cistarget/tf_lists/allTFs_mm.txt.

Gonadal phenotype annotation was constituted using the Mouse Phenotype OBO database v1.2 (Open Biological and Biomedical Ontology) from MGI (Mouse Genome Informatics) (https://www.informatics.jax.org/downloads/reports/Mpheno_OBO.ontology). Phenotype names containing the patterns “gonad*”, “testi*”, “ovar*”, “fertility”, and “sex*” were selected. Genes associated with the selected phenotype IDs were retrieved from the gene-phenotype database from MGI (https://www.informatics.jax.org/downloads/reports/MGI_GenePheno.rpt).

The list of the 1,860 mouse transcription factors and the 22,215 genes associated with a gonadal phenotype is provided in the **data file S3**.

### ATAC-seq, library prep and sequencing

Sorted Sertoli and pre-granulosa cells (∼60,000) were isolated from E11.5, E12.5, E13.5 and E15.5 gonads via FACS. ATAC-seq library preparation was performed using the Active Motif kit (Active Motif, cat. 53150) according to the manufacturer instructions. Sample concentration was measured using Qubit ds HS Assay Kit (Invitrogen, 2326054) and sample distribution size using Tapestation with high sensitivity D1000 tape (Agilent, 5067-5585). Samples were then pooled together accordingly into a 4nM library and sequenced with 37 bp paired end (PE) reads on the Illumina Next-Seq 500 platform at the Bar-Ilan Sequencing Unit with 25-40 million reads per sample.

### ATAC-seq mapping, peak calling and quality controls

FastQ files were processed using nf-core/atacseq pipeline v2.1.2 (*103*). Briefly, read quality controls were performed with FastQC. Sequencing adaptors were removed with Cutadapt. Reads were mapped on the mm10/GRCm38 reference genome from Gencode (M25) with BWA. Mapped reads were filtered to remove the unpaired reads, the mitochondrial reads, the duplicated reads, the multimapped reads, the fragments with insert size > 2 kb, and the reads mapping to the blacklisted regions (https://github.com/Boyle-Lab/Blacklist/blob/master/lists/mm10-blacklist.v2.bed.gz). Blacklist region file has been modified to allow peak calling on the Y chromosome to inspect the open chromatin regions around the *Sry* gene.

Peaks were called with MACS2 using the “narrow_peaks” parameter. For each condition, peaks with FDR<0.01 and found in at least two replicates were merged as consensus open chromatin regions. The obtained consensus regions were then combined and merged to constitute the set of non-overlapping open chromatin regions present in any conditions and was used for the downstream analysis. Read quantification was performed on the obtained open chromatin regions using featureCounts. Regions from which the maximum coverage is fewer than 50 reads across all samples were discarded from the read quantification matrix for the subsequent analysis.

Bigwig files normalized by million mapped reads from the nf-core/atacseq pipeline were corrected to be normalized by the size factors calculated from reads in peaks from DESeq2 in order to make the peak height reflecting the downstream analysis.

Consensus peak annotation for each condition was assessed using ChIPseeker (*104*). Sample correlation (Spearman) and PCA was performed using consensus region read counts normalized with DESeq2 with VST (Variance Stabilizing Transformation).

### Differential chromatin accessibility analysis

Differential chromatin accessibility analysis was performed similarly to the differential expression analysis. Sex differential accessibility analysis was performed on the filtered read quantification matrix using DESeq2 with ∼Sex_stage as model and using the “Wald” test, while stage-specific differential accessibility analysis was performed using the “LTR” test with ∼Stage as model. In both analyses, regions with log2(FoldChange) < −1 and log2(FoldChange) > 1, as well as an adjusted *p*-value < 0.01 were considered as differentially accessible.

Accessibility values (filtered read count matrix normalized by library size) from the stage differentially accessible regions were transformed as z-scores, clustered into groups according to their accessibility profiles using hclust and “ward.D2” method, and represented as heatmap using ComplexeHeatmap. The optimal number of clusters were assessed by visual inspection.

### Open chromatin region and gene expression correlations

Link between open chromatin region and gene expression was calculated using the same method used in the cisDynet R package (*105*). Briefly, for each protein-coding gene, we selected the open chromatin regions within 500 kb upstream and downstream of the canonical TSS of the gene as potential regulatory regions. Then we calculated the Pearson correlation coefficients between the gene expression and the chromatin accessibility of each region (all stages and sexes together). To control for false positives, we selected 10,000 random open chromatin regions and calculated the correlation coefficients between the regions and expression. We then tested the significance of each region-to-gene correlation using the *Z*-test method and obtained a *p*-value that is corrected for false discovery rate (FDR). Finally, we considered a significant region-to-gene link with a Pearson correlation coefficient[>0.7 (positive link) and <0.7 (negative link) and a FDR[<0.01. We excluded open chromatin regions overlapping TSS of alternative transcripts from our analysis to avoid links between genes and their alternative promoters. We visualized the region-to-gene links using Gviz and GenomicInteraction packages (*106*, *107*).

### Promoter Capture Hi-C

Capture Hi-C libraries (3 libraries of E13.5 *Sox9^IRES-GFP^* Sertoli cells and 2 libraries of E13.5 Enh8-mCherry pre-granulosa cells) were generated from 100,000 sorted cells, crosslinked as previously described (*108*, *109*). Following lysis, the nuclei were permeabilized and digested with DpnII (NEB) overnight. The restriction overhangs were filled in using a biotinylated dATP (Jena Bioscience) and ligation was performed for 4 hours at 16°C (T4 DNA ligase; Life Technologies). The crosslinks were reversed using proteinase K and overnight incubation at 65°C, followed by purification with SPRI beads (AMPure XP; Beckman Coulter). Short fragments up to 1000 bp were produced via tagmentation and the biotinylated restriction junctions were then pulled down using MyOne C1 streptavidin beads (Life Technologies). PCR amplification (5 cycles) was performed on the libraries directly bound to the C-1 beads and the libraries were purified using SPRI beads as before. Promoter Capture was performed using the mouse custom-designed Agilent SureSelect system, following the manufacturer’s protocol, followed by 7 PCR cycles. The libraries were sequenced using 150 bp paired-end sequencing on an Illumina NovaSeq X Plus (Novogene UK) with a sequencing depth of approximately 1bln paired reads for each biological replicate.

### PCHi-C data processing and detection of significant contacts

PCHi-C reads from biological replicates were merged using *cat* before processing further. Read processing, alignment and filtering was performed using a modified version of the Hi-C User Pipeline (HiCUP) v0.7.4, HiCUP Combinations (https://github.com/StevenWingett/HiCUP/tree/combinations), which first creates all possible combinations of ditags from the paired reads prior to mapping, then performs standard filtering for common Hi-C artefacts. To generate the consensus interaction calling per cell type (Granulosa and Sertoli cells), CHiCAGO input (.chinput) files were generated per biological replicate, using bam2chicago.sh from chicagoTools v1.13 and then CHiCAGO pipeline was run at 5 kb bins resolution taking the variation across replicates into account. The pipeline was run in two complementary modes. One where we only considered chromosomal interactions with the original baited fragment (termed “original bait”) and another where the baited fragment was expanded to a 5 kb bin resolution (termed “5 kb extended bait”), as we have previously described. We performed tests for epigenetic marker enrichment at promoter interacting regions in both modes, using the peakEnrichment4Features function in CHiCAGO. For the enrichment analysis ChIP-seq data for H3K4me3, H3K27ac and H3H27me3 from E13.5 pre-granulosa and Sertoli cells was obtained from GSE118755 and GSE130749 and reanalysed using the nf-core/chipseq v2.0.0 as previously described (*73*).

### TFBS motif analysis

Differential transcription factor motif enrichment analysis was performed with monaLisa (*110*) using vertebrate TFBS matrices of transcription factors from JAPSAR2024 (*111*) extracted using TFBSTools (*112*).

TFBS motifs differentially enriched in the sexually dimorphic accessible regions were analysed by merging the sexually dimorphic regions from all stages and by binning them by sex-specific accessibility. TFBS motifs differentially enriched in the dynamically accessible regions were analysed by binning the region by their cluster of dynamics profile.

Enrichment test was run against the bins as well as against a random genomic background (corrected for the GC content of the tested regions) to avoid GC content biases. TFBS motifs from transcription factors not expressed in the Sertoli and pre-granulosa cell samples (TPM<10) were filtered out.

TFBS motifs were clustered by enrichment score (log2(enrichment)) using hclust (“ward.D” method). For each enrichment cluster, TFBS motifs were grouped by similarity using motifStack (*113*) with cutoffPval = 0.001. The result is visualized as heatmaps of the log2(enrichment) showing the logos and the names of the merged TFBSs using ComplexeHeatmap (*114*).

### ATAC-seq footprint analysis

ATAC-seq footprint analysis was performed using TOBIAS v0.17.0 (*79*). We merged the bam files from the different replicates prior to the analysis. We ran ATACorrect and ScoreBigwig commands with default parameters. We then ran BINDetect on the combined sexually dimorphic regions from all stages and using TFBS motifs merged by similarity using motifStack as described above. Differential binding scores between pre-granulosa and Sertoli at each developmental stage cells were plot using ggplot2.

### Single-cell RNA-seq expression profiles

Single-cell RNA-seq data were obtained from GSE184708 (*9*). Data were loaded in Seurat v5.1.0 (*115*) using the gene, barcode, expression matrix and metadata files provided on GEO. Data was log-normalized and gene expression plotted using the VlnPlot function on selected cell types.

## Acknowledgements

We are grateful to the BIU Life Sciences Kanbar Core Facility unit of NGS, Flow Cytometry and Microscopy units. We are also grateful to the Animal Research Facility at BIU. We are thankful to the Genetic Modification Service of the Francis Crick Institute, and to the genotoul bioinformatics platform Toulouse Occitanie (Bioinfo Genotoul, https://doi.org/10.15454/1.5572369328961167E12) for providing the computing resources necessary for the current study. We thank members of our lab for advice, support, and helpful comments.

## Author contribution

Conceptualization, IS, EA, MR, LS, RLB and NG; Investigation, EA, MR, LS, CRF, DMM, VM, and NG; Methodology, IS, EA, MR, LS, RLB, and NG; Data curation, IS, LS, VM, and NG; Formal analysis, IS, LS, VM, and NG; Visualization, IS, EA, MR, RW, LS, VM, and NG; Software, IS, and RW; Resources, EA, MR, LS, RLB, VM, and NG; Writing – review & editing, IS, EA, MR, RLB, VM, and NG; Project administration, VM and NG; Supervision, RLB, VM, and NG; Funding acquisition, RLB, VM, and NG. All authors have read and accepted the data being presented in the manuscript.

## Competing declaration

The authors declare that they have no competing interests.

## Data and materials availability

All data is available in the manuscript or the supplementary materials. RNA-seq, ATAC-seq and PCHi-C data have been deposited in the Gene Expression Omnibus under accession number GSE277647 and GSE277646 and GSE293342, respectively. The code produced to analyze the data is available on Zenodo: https://doi.org/10.5281/zenodo.15222305.

