## Supplementary materials for "The gene regulatory landscape driving mouse gonadal supporting cell differentiation"

[4] The author is deceased.

[5] The Francis Crick Institute, 1 Midland Road, London NW1 1AT, UK.

[6] VIB Center for Molecular Neurology, VIB, Antwerp, Belgium.

[7] Department of Biomedical Sciences, University of Antwerp, Antwerp, Belgium.

[8] VIB Center for AI and Computational Biology, VIB, Leuven, Belgium.

### These authors contributed equally.

**Contact information:**

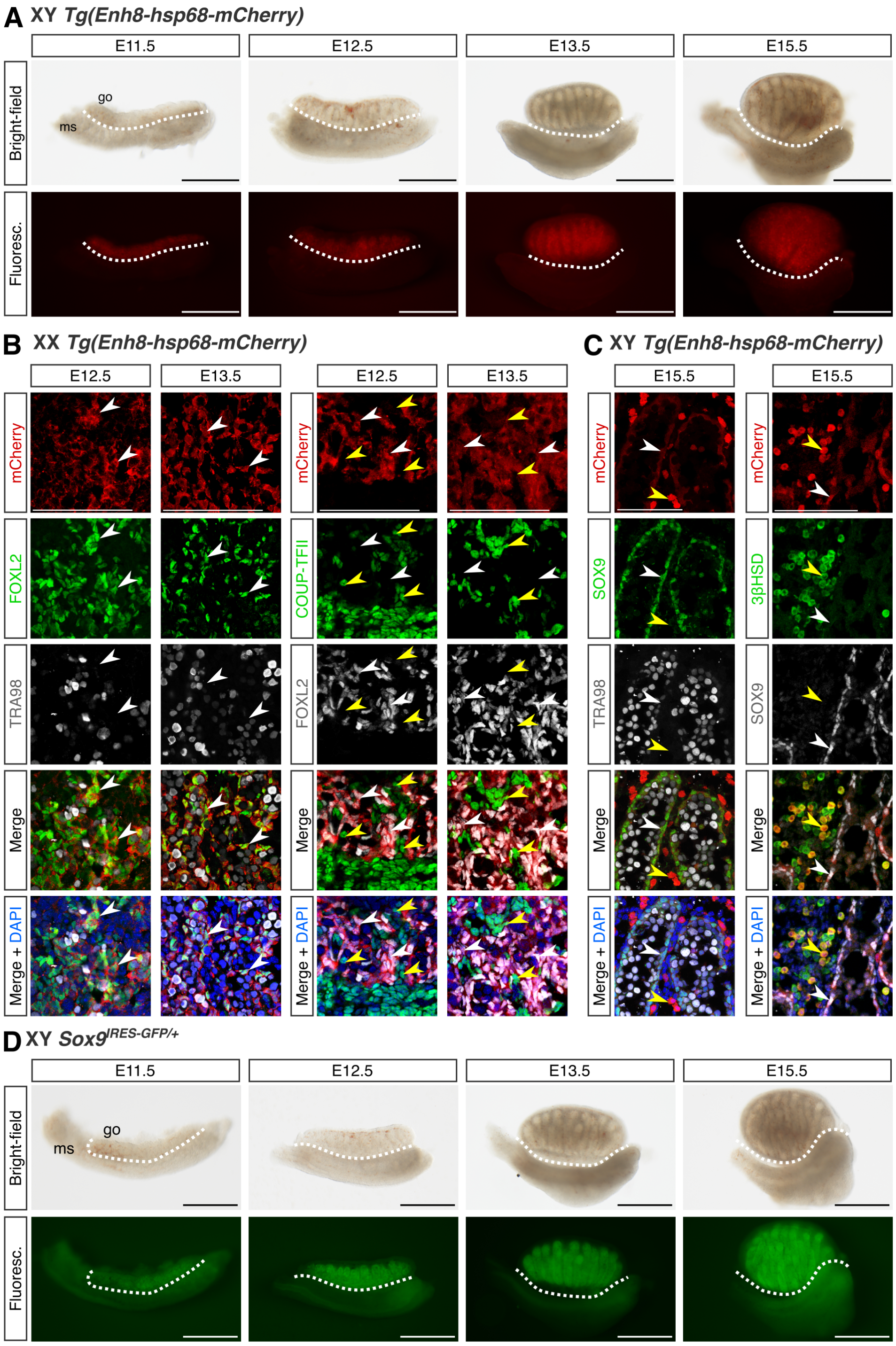


**Figure S1. Characterization of the *Tg(Enh8-hsp68-mCherry)* and *Sox9*$^{\text{IRES-GFP}}$ expressing cells in mouse fetal gonads.**

(**A**) Binocular pictures of dissected XY *Enh8-mCherry* gonads at E11.5, E12.5, E13.5 and E15.5 in bright-field and fluorescence. go=gonad, ms=mesonephros. (**B**) Immunofluorescent staining of E12.5 and E13.5 *Enh8-mCherry* ovary sections. In the two panels, mCherry (transgene) is labelled in red. In the left hand side panel, FOXL2, a marker of pre-granulosa cells is labelled in green, and TRA98, a marker of germ cells is labelled in grey. In the right hand side, COUP-TFII (also known as NR2F2), marker of stromal progenitor cells, is labelled in green, and FOXL2 in grey. The white arrowheads point mCherry-FOXL2 positive cells, while the yellow arrowheads show cells only positive for COUP-TFII. Scale bar: 100 µm. (**C**) Immunofluorescent staining of E13.5 *Enh8-mCherry* testis sections. On the first column, mCherry (transgene) is labelled in red, SOX9, a marker of Sertoli cells, is labelled in green, and TRA98, a marker of germ cells is labelled in grey. On the second column, mCherry (transgene) is labelled in red, 3BHSD, a marker of Leydig cells, are labelled in green, and SOX9 is labelled in grey. The white arrowheads point mCherry-SOX9 positive cells, while the yellow arrowheads show interstitial mCherry cells positive for 3BHSD. Scale bar: 100 µm. (**D**) Binocular pictures of dissected XY *Sox9*$^{\text{IRES-GFP}}$ gonads at E11.5, E12.5, E13.5 and E15.5 in bright-field and fluorescence.

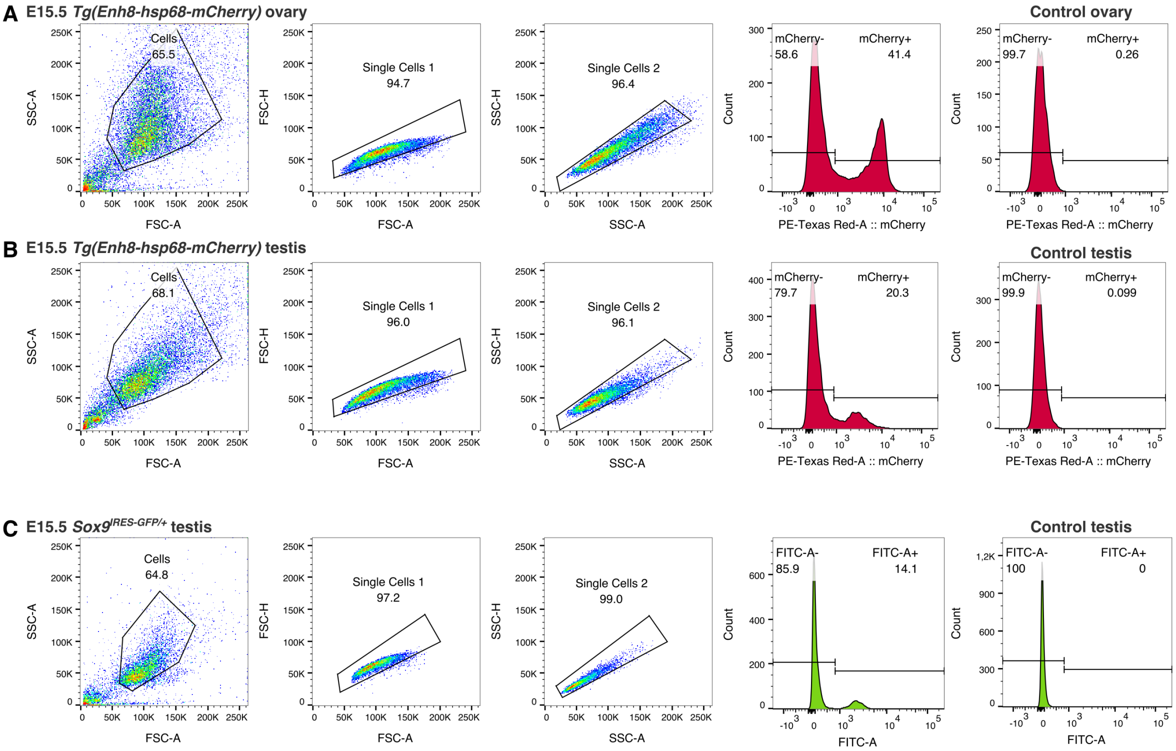


**D**

| Sex | Type of mice | Developmental stage | Supporting cell ratio (%) |
|---|---|---|---|
| XY | Nr5a1-GFP, whole gonads* *(CD1 background)* | E11.5 | 18** |
| | | E12.5 | 17 |
| | | E13.5 | 15 |
| | | E16.5 | 22 |
| | *TESCO-CFP*⁺ cells *(C57BL6/J x CBA-Ca F1 background)* | E13.5 | 7 |
| | | E15.5 | 5 |
| | *SOX9*$^{IRES\text{-}GFP+}$ cells *(C57BL6/J x CBA-Ca F1 background)* | E13.5 | 14 |
| | | E15.5 | 15 |
| XX | Nr5a1-GFP, whole gonads* *(CD1 background)* | E11.5 | 17** |
| | | E12.5 | 35 |
| | | E13.5 | 37 |
| | | E16.5 | 35 |
| | *TESMS-CFP*⁺ cells *(C57BL6/J x CBA-Ca F1 background)* | E13.5 | 6 |
| | | E15.5 | 6 |
| | *Enh8-mCherry*⁺ cells *(C57BL6/J x CBA-Ca F1 background)* | E13.5 | 33 |
| | | E15.5 | 35 |

* Based on Mayère et al. scRNA-seq *(9)*
** Gonad-mesonephros

**Figure S2. Cell sorting of the *Enh8-mCherry* and *Sox9*$^{IRES\text{-}GFP}$ expressing cells from mouse fetal gonads.**

**(A)** and (**B**) FACS gating of the ovarian and testicular mCherry positive cells at E15.5 gonads, respectively. First, cells are gated according to their size (FSC) and their light refraction (SSC). Second, cell doublets are filtered out (FSC-A vs FSC-H), and finally, mCherry positive cells are selected using their fluorescence level (PE-Texas Red-A) compared to a control gonad (shown on the right-hand side). **(C)** FACS gating of the testicular GFP positive cells from E15.5 testes. Gating was made similarly to the mCherry cells, but using FITC to measure the GFP signal. **(D)** Table recapitulating the percentage of supporting cells in wild type fetal gonads (taken from scRNA-seq data, Mayère et al. *(9)*) and obtained by FACS using previously used transgenes, *TESCO-CFP* and *TESMS-CFP*, and using the current *Sox9*$^{IRES\text{-}GFP}$ and *Enh8-mCherry*, that label Sertoli and pre-granulosa cells, respectively. The genetic background of each mouse lines is indicated.

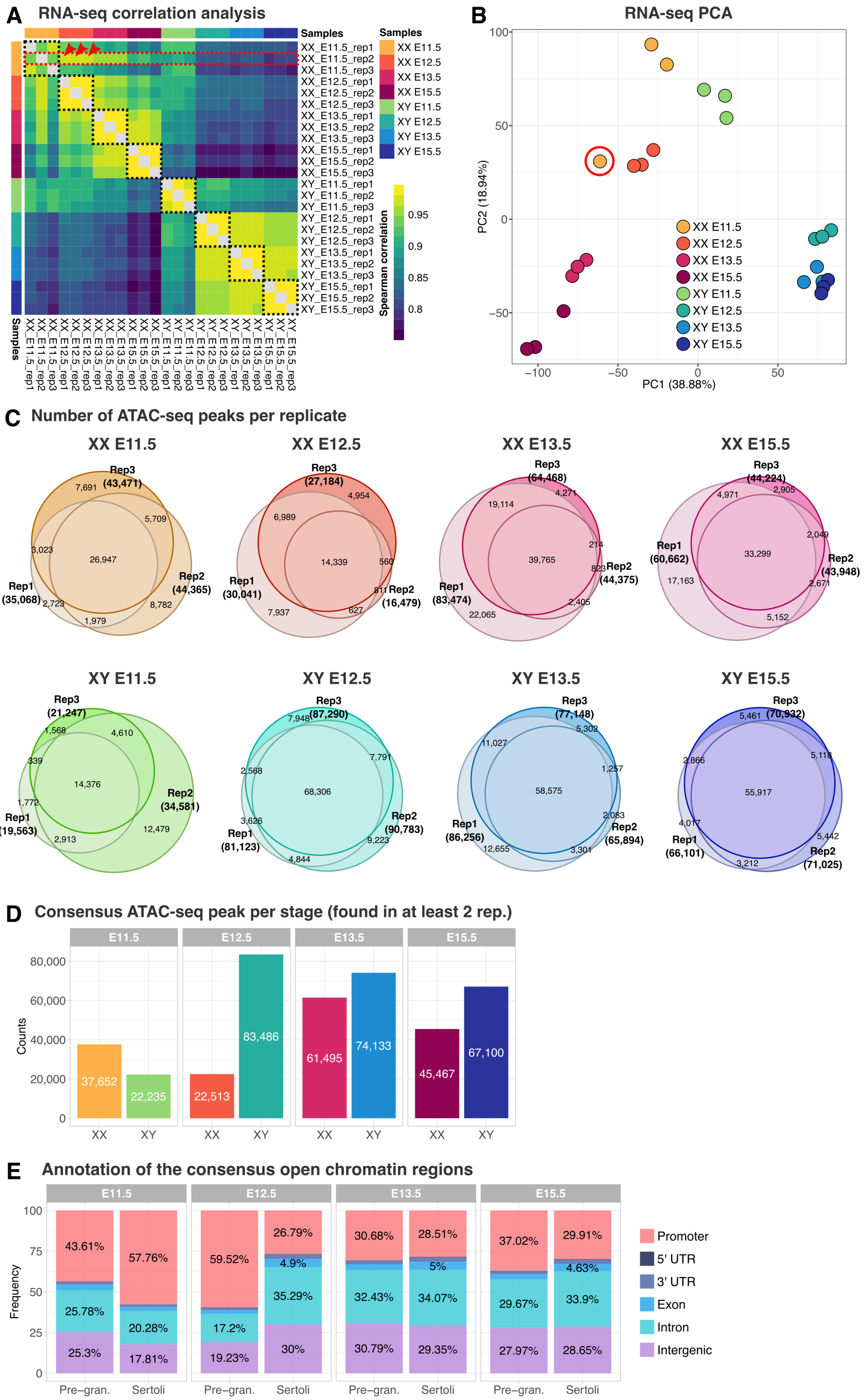
A
RNA-seq correlation analysis
Samples
XX_E11.5_rep1
XX_E11.5_rep2
XX_E11.5_rep3
XX_E12.5_rep1
XX_E12.5_rep2
XX_E12.5_rep3
XX_E13.5_rep1
XX_E13.5_rep2
XX_E13.5_rep3
XX_E15.5_rep1
XX_E15.5_rep2
XX_E15.5_rep3
XY_E11.5_rep1
XY_E11.5_rep2
XY_E11.5_rep3
XY_E12.5_rep1
XY_E12.5_rep2
XY_E12.5_rep3
XY_E13.5_rep1
XY_E13.5_rep2
XY_E13.5_rep3
XY_E15.5_rep1
XY_E15.5_rep2
XY_E15.5_rep3
XX E11.5
XX E12.5
XX E13.5
XX E15.5
XY E11.5
XY E12.5
XY E13.5
XY E15.5
Spearman correlation
0.95
0.9
0.85
0.8
B
RNA-seq PCA
PC1 (38.88%)
PC2 (18.94%)
100
50
0
−50
−100
C
Number of ATAC-seq peaks per replicate
XX E11.5
Rep3 (43,471)
7,691
5,709
3,023
26,947
Rep1 (35,068)
2,723
1,979
8,782
Rep2 (44,365)
XX E12.5
Rep3 (27,184)
4,954
6,989
14,339
560
Rep1 (30,041)
611
Rep2 (16,479)
7,937
627
XX E13.5
Rep3 (64,468)
4,271
19,114
214
39,765
823
Rep2 (44,375)
Rep1 (83,474)
2,405
22,065
XX E15.5
Rep3 (44,224)
4,971
2,905
2,049
Rep1 (60,662)
33,299
Rep2 (43,948)
17,163
2,671
5,152
XY E11.5
Rep3 (21,247)
1,568
4,610
339
14,376
Rep2 (34,581)
1,772
Rep1 (19,563)
2,913
12,479
XY E12.5
Rep3 (87,290)
7,948
7,791
2,568
68,306
3,626
Rep2 (90,783)
Rep1 (81,123)
9,223
4,844
XY E13.5
Rep3 (77,148)
5,302
11,027
1,257
58,575
2,083
Rep1 (86,256)
Rep2 (65,894)
12,655
3,301
XY E15.5
Rep3 (70,932)
5,461
5,118
2,866
55,917
4,017
5,442
Rep1 (66,101)
Rep2 (71,025)
3,212
D
Consensus ATAC-seq peak per stage (found in at least 2 rep.)
E11.5
E12.5
E13.5
E15.5
Counts
80,000
60,000
40,000
20,000
0
37,652
22,235
22,513
83,486
61,495
74,133
45,467
67,100
XX
XY
E
Annotation of the consensus open chromatin regions
Frequency
100
75
50
25
0
E11.5
43.61%
25.78%
25.3%
57.76%
20.28%
17.81%
E12.5
59.52%
17.2%
19.23%
26.79%
4.9%
35.29%
30%
E13.5
30.68%
32.43%
30.79%
28.51%
5%
34.07%
29.35%
E15.5
37.02%
29.67%
27.97%
29.91%
4.63%
33.9%
28.65%
Pre−gran.
Sertoli
Promoter
5' UTR
3' UTR
Exon
Intron
Intergenic

**Figure S3. Quality controls of the RNA-seq and the ATAC-seq data.**

(**A**) Correlation matrix of the RNA-seq samples calculated using the normalized read counts. Correlations between sample replicates are surrounded with black dashed lines. Sample replicates correlate strongly between each other except XX E11.5 replicate 2, highlighted by red dashed lines, that correlates more with the XX E12.5 samples rather that the other XX E11.5 replicates as shown by the red arrowheads. (**B**) PCA of the RNA-seq samples calculated using the normalized read counts. XX E11.5 replicate 2 is surrounded by a read circle. XX E11.5 replicate 2 is closer to the XX E12.5 than the other XX E11.5 replicates. This sample was considered as an outlier and was removed from the downstream analysis. (**C**) Venn diagrams showing the number of unfiltered ATAC-seq peaks called in each sample and their overlap between the biological replicates. (**D**) Number of filtered ATAC-seq peaks detected in at least two replicates, so called consensus peaks, for each condition. (**E**) Percentage of genomic features overlapped by the consensus ATAC-seq peaks for each condition.

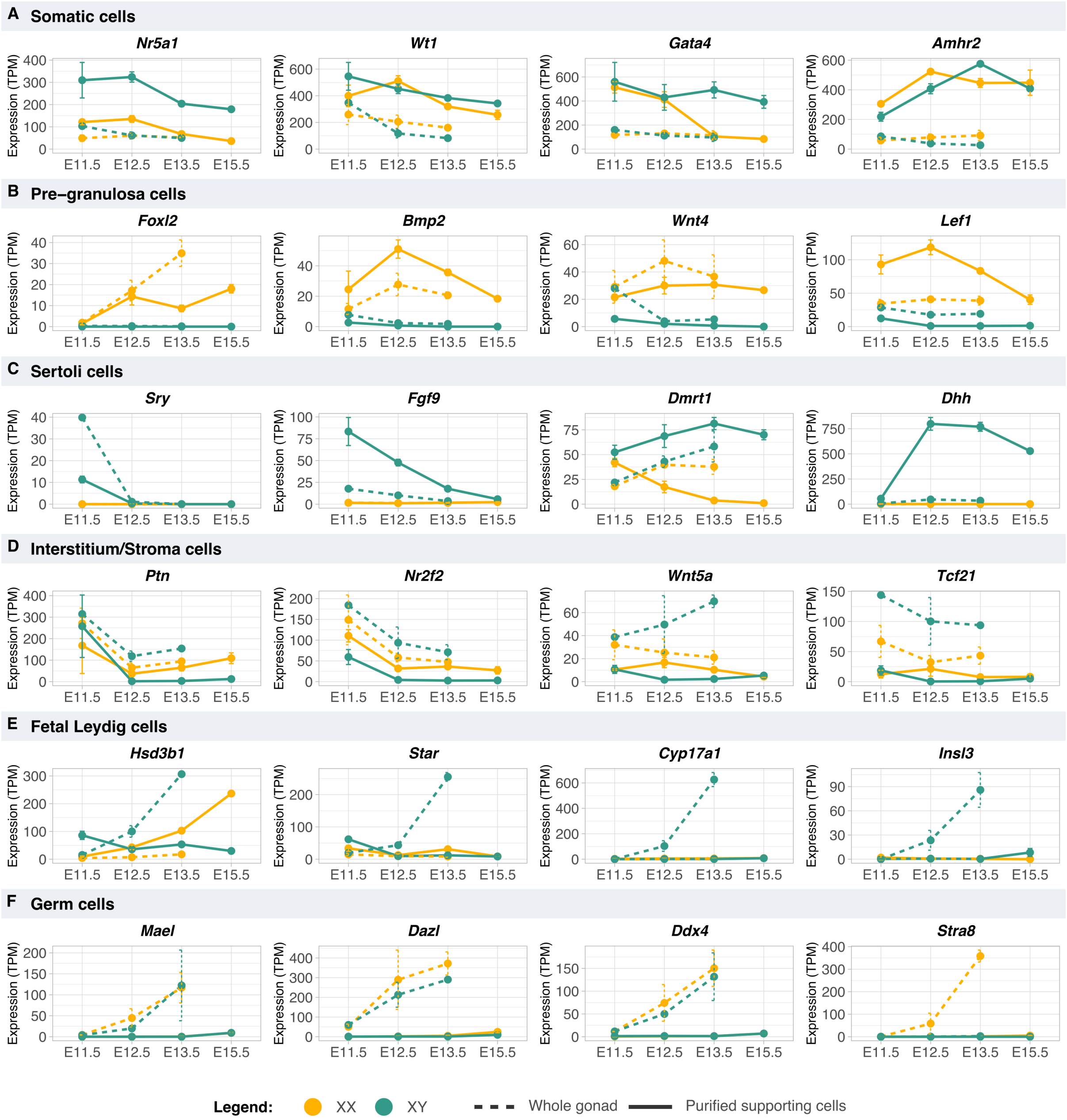


**Figure S4. Expression of marker genes for the most abundant cell types of the gonad.**

(**A-F**) Expression profiles in TPM of marker genes in purified Sertoli and pre-granulosa cells (continuous line) and in whole gonads (from Zhao et al. *(58)*, dashed line) along embryonic stages in both sexes (XX in yellow, XY in green).

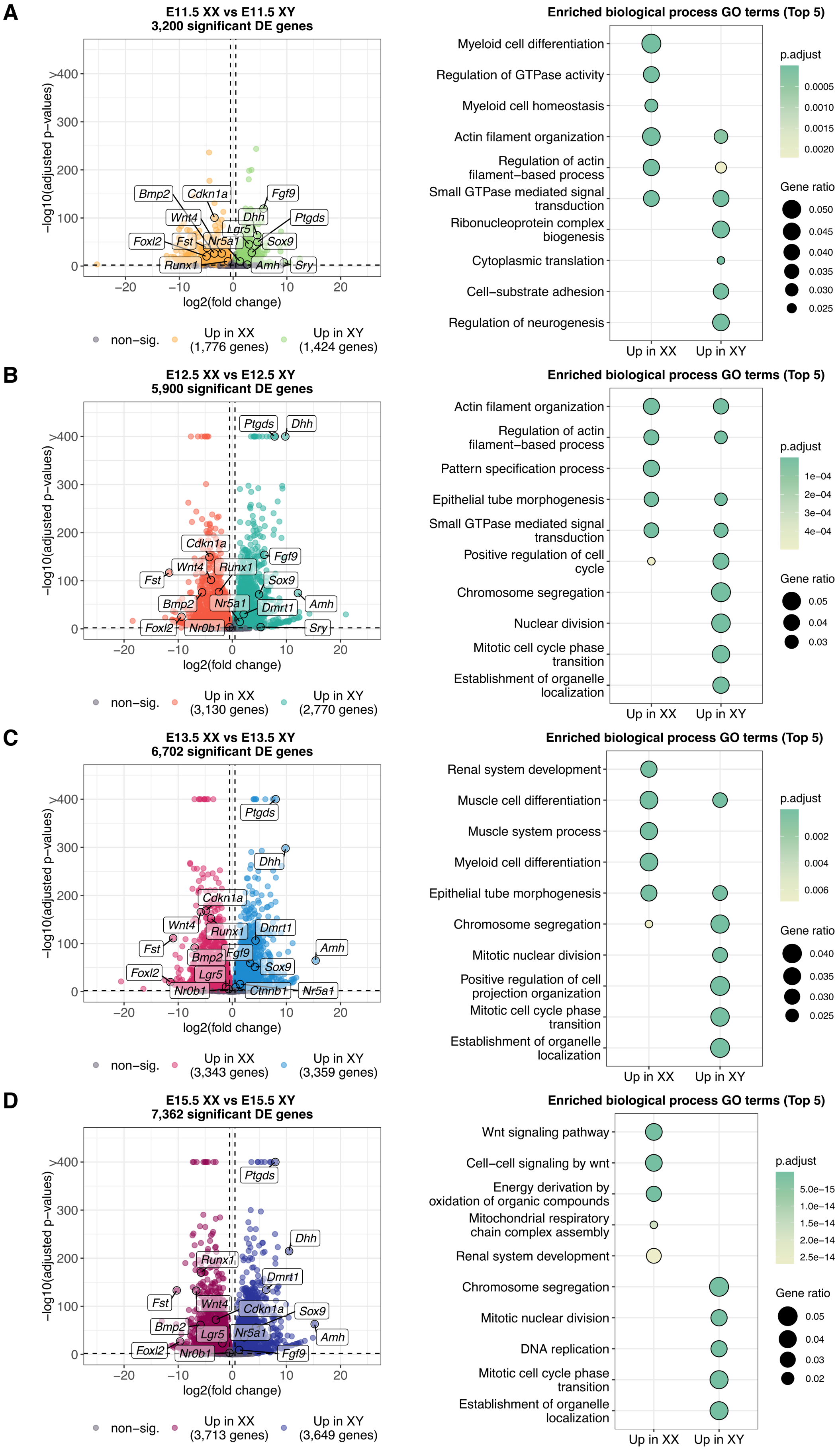
A
E11.5 XX vs E11.5 XY
3,200 significant DE genes
−log10(adjusted p−values)
log2(fold change)
Bmp2
Cdkn1a
Fgf9
Wnt4
Dhh
Ptgds
Lgr5
Foxl2
Fst
Nr5a1
Sox9
Runx1
Amh
Sry
non−sig.
Up in XX (1,776 genes)
Up in XY (1,424 genes)
Enriched biological process GO terms (Top 5)
Myeloid cell differentiation
Regulation of GTPase activity
Myeloid cell homeostasis
Actin filament organization
Regulation of actin filament−based process
Small GTPase mediated signal transduction
Ribonucleoprotein complex biogenesis
Cytoplasmic translation
Cell−substrate adhesion
Regulation of neurogenesis
Up in XX
Up in XY
p.adjust
Gene ratio
B
E12.5 XX vs E12.5 XY
5,900 significant DE genes
Ptgds
Dhh
Cdkn1a
Fgf9
Wnt4
Runx1
Fst
Sox9
Amh
Bmp2
Nr5a1
Dmrt1
Foxl2
Nr0b1
Sry
Up in XX (3,130 genes)
Up in XY (2,770 genes)
Actin filament organization
Regulation of actin filament−based process
Pattern specification process
Epithelial tube morphogenesis
Small GTPase mediated signal transduction
Positive regulation of cell cycle
Chromosome segregation
Nuclear division
Mitotic cell cycle phase transition
Establishment of organelle localization
C
E13.5 XX vs E13.5 XY
6,702 significant DE genes
Ptgds
Dhh
Cdkn1a
Wnt4
Runx1
Dmrt1
Fst
Amh
Bmp2
Fgf9
Sox9
Foxl2
Lgr5
Nr0b1
Ctnnb1
Nr5a1
Up in XX (3,343 genes)
Up in XY (3,359 genes)
Renal system development
Muscle cell differentiation
Muscle system process
Myeloid cell differentiation
Epithelial tube morphogenesis
Chromosome segregation
Mitotic nuclear division
Positive regulation of cell projection organization
Mitotic cell cycle phase transition
Establishment of organelle localization
D
E15.5 XX vs E15.5 XY
7,362 significant DE genes
Ptgds
Dhh
Runx1
Dmrt1
Fst
Wnt4
Cdkn1a
Sox9
Bmp2
Lgr5
Nr5a1
Amh
Foxl2
Nr0b1
Fgf9
Up in XX (3,713 genes)
Up in XY (3,649 genes)
Wnt signaling pathway
Cell−cell signaling by wnt
Energy derivation by oxidation of organic compounds
Mitochondrial respiratory chain complex assembly
Renal system development
Chromosome segregation
Mitotic nuclear division
DNA replication
Mitotic cell cycle phase transition
Establishment of organelle localization

**Figure S5. Differentially expressed genes between pre-granulosa and Sertoli cells from E11.5 to E15.5.**

(**A-D**) Left-hand side plots are volcano plots showing the differentially expressed genes at each developmental stage. Known gonadal factors are indicated. Right-hand side plots represent the top 5 GO terms of the pre-granulosa and Sertoli cell differentially expressed genes.

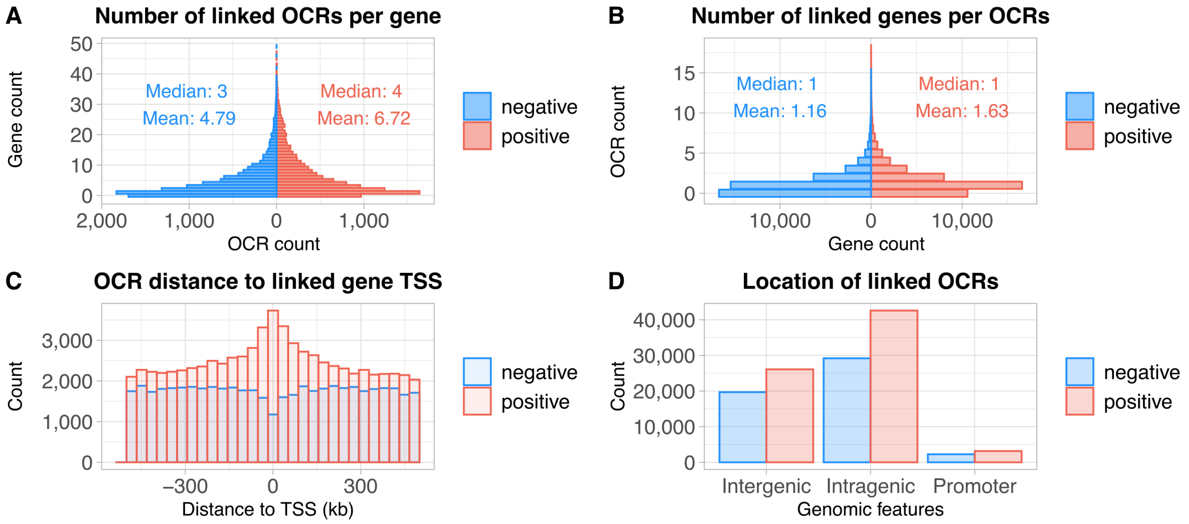


**Figure S6. Characterization of the open chromatin regions that correlate positively and negatively with gene expression.**

(**A**) Distribution of the number of genes linked with open chromatin regions (OCRs) with a positive (red) or negative (blue) correlation. (**B**) Distribution of the number of OCRs linked with genes with a positive (red) or negative (blue) correlation. (**C**) Distribution of the distance of OCRs to the TSS of their linked genes. (**D**) Genomic features overlapped by the linked OCRs presenting linkage to target genes.

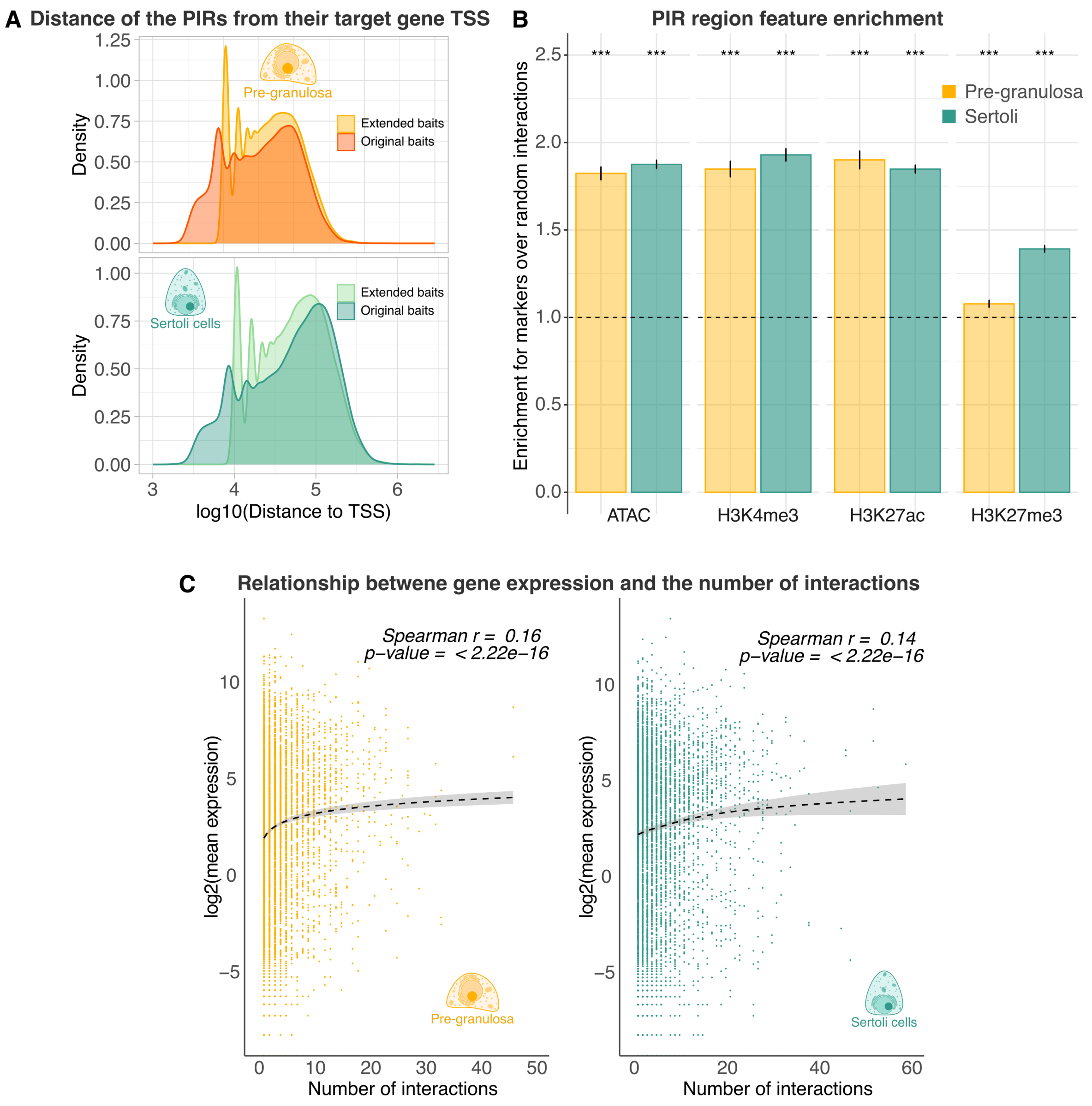


**Figure S7. Promoter-capture Hi-C analysis.**

(**A**) Distance of the promoter-interacting regions (PIRs) from their target genes (or bait) in the two analysis modes with (5 kb extended baits) and without (original baits) inclusion of promoters in the binning process in pre-granulosa and Sertoli cells. (**B**) Enrichment of the PIRs in different chromatin features in pre-granulosa and Sertoli cells. ATAC-seq data corresponds to our E13.5 data, the histone marks are taken from Garcia-Moreno et al. *(41)* and analysed as in (73). (**C**) Correlation between gene expression and the number of PIR per gene in pre-granulosa and Sertoli cells. RNA-seq data corresponds to our E13.5 data.

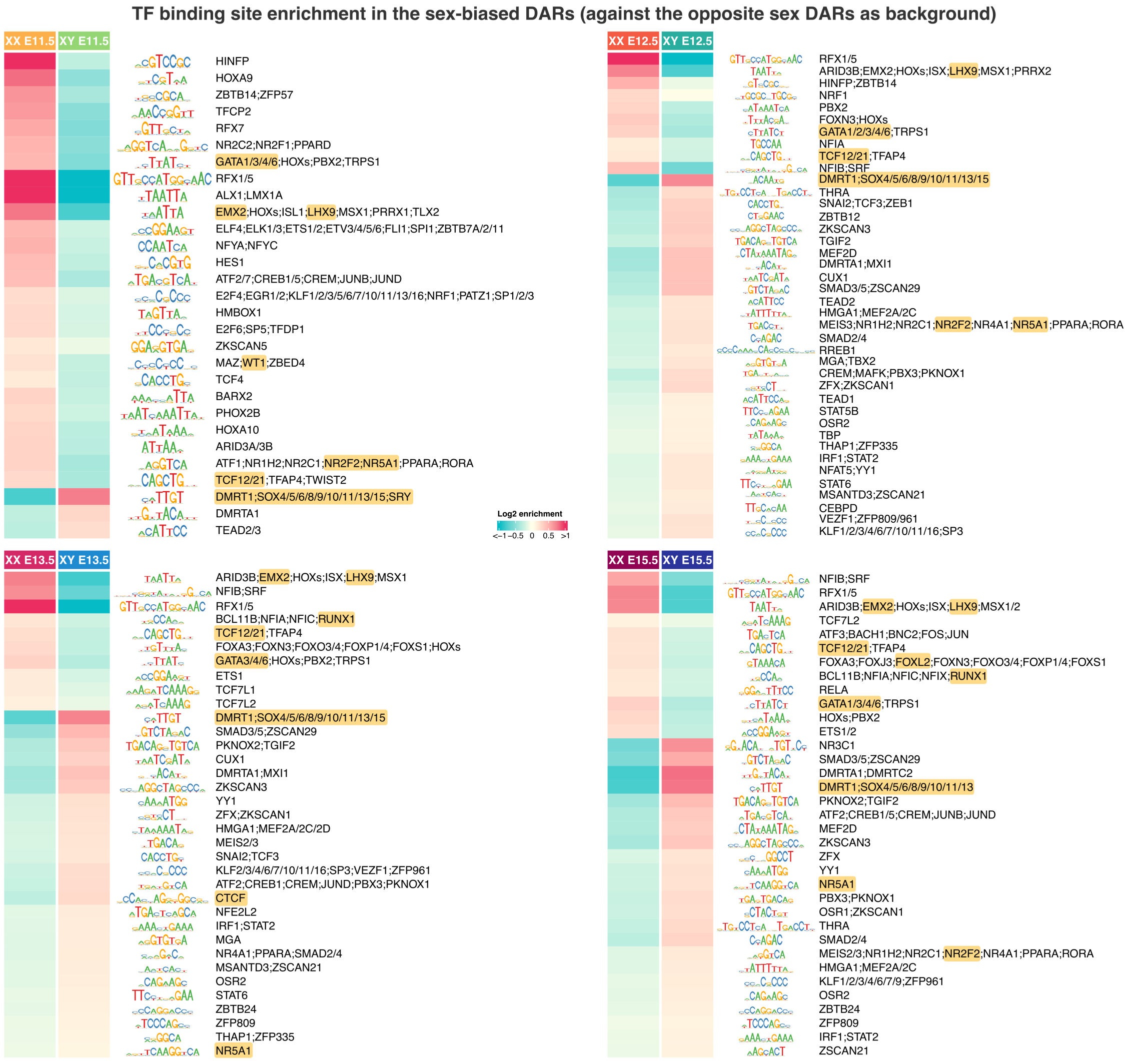


**Figure S8. Differential transcription factor motif enrichment in the sexually dimorphic open chromatin regions compared to the opposite sex regions.**

Heatmaps showing the enrichment score of expressed transcription factor motifs in pre-granulosa and Sertoli cell differentially accessible regions at each developmental stage. Motifs were merged by similarity and the consensus logo and the corresponding transcription factors are indicated on the right-hand side. Noticeable transcription factors are highlighted in yellow.

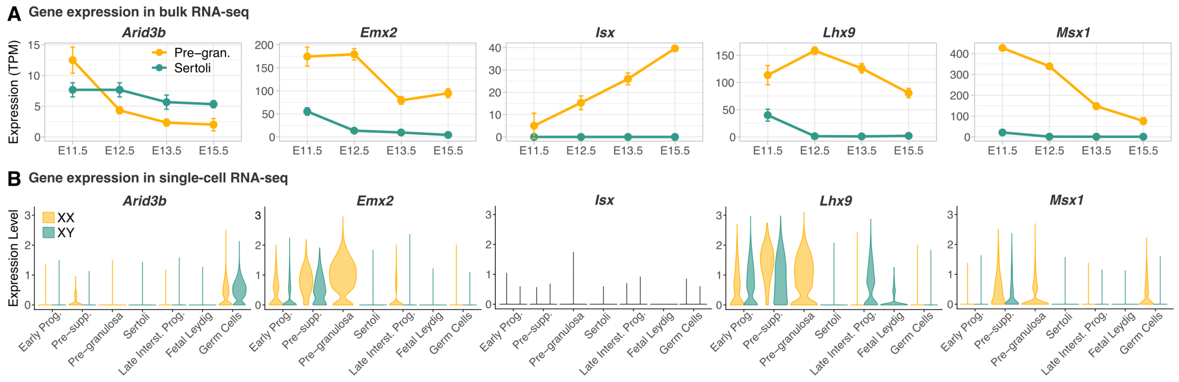


**Figure S9. Expression profiles of TFs in bulk supporting cell RNA-seq and whole gonad scRNA-seq.**

(**A**) and (**B**) Expression profiles of the TFs the most differentially bound in the pre-granulosa cell-biased open chromatin regions in either supporting cell bulk RNA-seq or in the gonadal single-cell RNA-seq atlas (log-normalized expression) *(9)*, respectively.

**Table S1. Primers for genotyping mice and cloning**

| Target | Primer name | Description | Sequence 5' to 3' | Product and size |
|---|---|---|---|---|
| Sex | Sex_F | X/Y chromosome | 5'-GATGATTTGAGTGGAAATGTGAGGTA-3' | Sex_F + Sex_R: 280bp in XY; 685bp,660bp and 480bp in XX McFarlane et al. *(97)* |
| | Sex_R | X/Y chromosome | 5'-CTTATGTTTATAGGCATGCACCATGTA-3' | |
| mCherry | mCherry_F | mCherry transgene | 5'–GATAACATGGCCATCATCAAG–3' | mCherry_F + mCherry_R: 248 bp in positive transgene |
| | mCherry_R | mCherry transgene | 5'– CTTCAAGTAGTCGGGGATG- 3' | |
| GFP (for X-GFP or Sox9-IRES-GFP mice) | GFP_F | GFP transgene | 5'-TGCAGTGCTTCAGCCGCTAC-3' | GFP_F + GFP_R: 400 bp in positive transgene |
| | GFP_R | GFP transgene | 5'-CCAGCAGGACCATGTGATCG-3' | |
| GFP homozygousity (for Sox9-IRES GFP het/hom) | GFP_hom_F | Flanks the 3'-UTR of Sox9 | 5'-GGCTTGTCTCCTTCAGAG-3' | GFP_hom_F + GFP_hom_R + IRES_R: 120 bp in WT allele and 450 bp in KI allele |
| | GFP_hom_R | Flanks the 3'-UTR of Sox9 | 5'-GAG AAT ATT CCT CAC AGA GG-3' | |
| | IRES_R | Flanks the IRES sequence | 5'-CTCTTTCCACCAACTATCC-3' | |
| Enh8-hsp68-mCherry | Enh8-AseI_F | Amplifies the Enh8 along with AseI restriction site | 5' CGCCATGCATTAGTTATTAATTCTTAAAA CAGCACCCACCAC-3' | Used to clone Enh8 into the hsp68-mCherry vector into the AseI site |
| | Enh8_AseI_R | Amplifies the Enh8 along with AseI restriction site | 5' CGGTACGYAATCAAATTAATTTTCCTGGG TGATATAGTATGC-3' | |

#### Table S2. Antibodies/ Dyes used in this study

| Target | Host | Manufacture | Cat. number | Dilution | Antibody type | Used for staining | Cell type |
|---|---|---|---|---|---|---|---|
| αSOX9 | Goat | R&D systems | AF3075 | 1:300 | Primary | Embryonic gonads | Sertoli cells |
| αFOXL2 | Rabbit | Abcam | ab246511 | 1:250 | Primary | Embryonic gonads | Pre-granulosa cells |
| αCOUP-TFII | Mouse | R&D systems | PP-H7147-00 | 1:200 | Primary | Embryonic gonads | Stromal progenitor cells |
| αCHERRY | Goat | Rockland | 200-101-379 | 1:100 | Primary | Embryonic gonads | CHERRY |
| αCHERRY | Rabbit | Rockland | 600-401-379 | 1:200 | Primary | Embryonic gonads | CHERRY |
| αGCNA1 (TRA98) | Rat | Abcam | ab82527 | 1:200 | Primary | Embryonic gonads | Germ cells |
| αRabbit-Alexa flour 488 | Donkey | Invitrogen | A-21206 | 1:500 | Secondary | Embryonic gonads | - |
| αGoat-Alexa flour 488 | Donkey | Invitrogen | A-11055 | 1:500 | Secondary | Embryonic gonads | - |
| αGoat-Alexa flour 568 | Donkey | Invitrogen | A-11057 | 1:500 | Secondary | Embryonic gonads | - |
| αRabbit-Alexa flour 568 | Donkey | Invitrogen | A-10042 | 1:500 | Secondary | Embryonic gonads | - |
| αRat-Alexa flour 647 | Donkey | Abcam | ab150155 | 1:500 | Secondary | Embryonic gonads | - |
| DAPI | - | Invitrogen | D-1306 | 300nM | Secondary | All | Nucleic acid |

**Data S1.**

RNA differential expression analysis of the genes exhibiting sexual dimorphism at each embryonic stage.

**Data S2.**

Lists of the mouse transcription factor encoding genes and the genes with known gonadal associated phenotypes extracted from the MGI database.

**Data S3.**

Differential expression analysis of the genes exhibiting expression changes in pre-granulosa and Sertoli cells along cell differentiation.

**Data S4.**

ATAC differential accessibility analysis of the regions exhibiting sexual dimorphism at each embryonic stage.

**Data S5.**

TF-binding motif enrichment in the sexually dimorphic accessible regions.

**Data S6.**

Differential accessibility analysis of the regions exhibiting accessibility changes in pre-granulosa and Sertoli cells along cell differentiation.

**Data S7.**

Differential TF-binding motif enrichment in the dynamically accessible regions.

**Data S8.**

Putative cis-regulatory element associated with their target genes with the linkage analysis.

**Data S9.**

Promoter-interacting regions from the PCHi-C analysis.

**Data S10.**

Gonadal TF footprint differential analysis and genomic location in pre-granulosa and Sertoli cells.
